# Targeted Down Regulation Of Core Mitochondrial Genes During SARS-CoV-2 Infection

**DOI:** 10.1101/2022.02.19.481089

**Authors:** Joseph W. Guarnieri, Joseph M. Dybas, Hossein Fazelinia, Man S. Kim, Justin Frere, Yuanchao Zhang, Yentli Soto Albrecht, Deborah G. Murdock, Alessia Angelin, Larry N. Singh, Scott L. Weiss, Sonja M. Best, Marie T. Lott, Henry Cope, Viktorija Zaksas, Amanda Saravia-Butler, Cem Meydan, Jonathan Foox, Christopher Mozsary, Yared H. Kidane, Waldemar Priebe, Mark R. Emmett, Robert Meller, Urminder Singh, Yaron Bram, Benjamin R. tenOever, Mark T. Heise, Nathaniel J. Moorman, Emily A. Madden, Sharon A. Taft-Benz, Elizabeth J. Anderson, Wes A. Sanders, Rebekah J. Dickmander, Victoria K. Baxter, Stephen B. Baylin, Eve Syrkin Wurtele, Pedro M. Moraes-Vieira, Deanne Taylor, Christopher E. Mason, Jonathan C. Schisler, Robert E. Schwartz, Afshin Beheshti, Douglas C. Wallace

**Affiliations:** The Children’s Hospital of Philadelphia, Philadelphia, PA 19104 USA; COVID-19 International Research Team; Kyung Hee University Hospital at Gangdong, Kyung Hee University, Seoul, South Korea; New York University, New York, NY 10016, USA; Rocky Mountain Laboratories NIAID, Hamilton, MT 59840; University of Nottingham, Nottingham, UK; University of Chicago, Chicago, IL, 60615, USA; Logyx, LLC, Mountain View, CA 94043, USA; NASA Ames Research Center, Moffett Field, CA 94035, USA; Weill Cornell Medicine, NY, 10065, USA; Scottish Rite for Children, Dallas, TX 75219, USA; University of Texas MD Anderson Cancer Center, Houston, TX, 77030, USA; University of Texas Medical Branch, Galveston, TX 77555, USA; Morehouse School of Medicine, Atlanta, GA 30310, USA; Iowa State University, Ames, IA 50011, USA; University of North Carolina, Chapel Hill, Chapel Hill, NC, 27599, USA; Johns Hopkins School of Medicine, Baltimore, MD 21287, USA; IB.University of Campinas, SP, Brazil; New York Genome Center, NY, USA; Broad Institute of MIT and Harvard, Cambridge, MA, 02142, USA; KBR, NASA Ames Research Center, Moffett Field, CA, 94035, USA; University of Pennsylvania, Philadelphia, PA 19104 USA

## Abstract

Defects in mitochondrial oxidative phosphorylation (OXPHOS) have been reported in COVID-19 patients, but the timing and organs affected vary among reports. Here, we reveal the dynamics of COVID-19 through transcription profiles in nasopharyngeal and autopsy samples from patients and infected rodent models. While mitochondrial bioenergetics is repressed in the viral nasopharyngeal portal of entry, it is up regulated in autopsy lung tissues from deceased patients. In most disease stages and organs, discrete OXPHOS functions are blocked by the virus, and this is countered by the host broadly up regulating unblocked OXPHOS functions. No such rebound is seen in autopsy heart, results in severe repression of genes across all OXPHOS modules. Hence, targeted enhancement of mitochondrial gene expression may mitigate the pathogenesis of COVID-19.

**One-Sentence Summary:** Covid-19 is associated with targeted inhibition of mitochondrial gene transcription.

---

COVID-19 is currently responsible for more than 6 million deaths and has infected close to 375 million people. While generally considered an inflammatory disease, recent evidence suggests that SARS-CoV-2 infection also inhibits mitochondrial bioenergetics, and that mitochondrial inhibition can activate of the inflammasome. Thus, in infected individuals, mitochondrial inhibition may not only contribute to excessive cytokine production but may be particularly impactful for the heart and brain, since these organs are highly reliant on mitochondrial energy production (*1–5*).

## SARS-CoV-2 Infection Modulates Mitochondrial Bioenergetics

SARS-CoV-2 infection has been found to markedly alter mitochondrial morphology in multiple human and monkey organs and cell lines (*6–8*). Infected Calu-3 cells showed mitochondria matrix condensation, swollen cristae, and decreased mitochondrial ATP synthase (complex V) subunit 5B (*7*). Lung-related microvascular endothelial cells (HULEC) and Human Pulmonary Alveolar Epithelial Cells (HPAEpiC) infected with SARS-CoV-2 demonstrated fragmented mitochondria with swollen cristae, increased mitochondrial reactive oxygen species (mROS) production, decreased oxidative phosphorylation (OXPHOS) complex I subunit NDUFA4, and decreased mitochondrial inner membrane protein TIMM23 (*6*). SARS-CoV-2 infection decreased both protein and transcript levels of the OXPHOS complex IV assembly factors COA4 and COA6 and the inner mitochondrial membrane protein TIMM8B, but increased glycolysis proteins including hexokinase (HK) isoforms 1, 2, and 3, and tricarboxylic acid (TCA) cycle malic enzymes 2 and 3 (*6, 9, 10*) **(Fig. S1)**.

Shifting the carbon source from glucose to galactose inhibited SARS-CoV-2 replication in TMPRSS-Vero E6 monkey and A549-hACE2 human cells (*11*). Treatment of CaCo-2 and Calu-3 cells with the HK inhibitor 2-deoxy-D-glucose (2-DG) suppressed viral replication (*9, 12*). Infected A549-hACE2 cells, human iPSC-derived airway organoids (hPSC-AO), and BALF and lung autopsy samples showed increased transcription of both HIF-1α and HIF-1α target genes involved in glycolytic metabolism (*10, 13, 14*). Treatment of hPSC-AO, human monocytes, and CaCo-2 cells with the HIF-1α inhibitors, chetomin or BAY87-2243, inhibited viral replication (*14–16*), and expressing inhibitory RNAs specific for HIF-1α in infected hPSC-AOs reduced viral replication, transcription of HIF-1α, and glycolysis gene expression (*14*).

SARS-CoV-2 infected human monocytes exhibited elevated glycolysis, suppressed OXPHOS, induced mROS production, increased hypoxia inducing factor 1-α (HIF-1α), up regulated HIF-1α target genes, increased viral load, and mRNA expression of IL-1β, IL-6, TNF and interferons. Treatment with 2-DG reduced viral load; and treatment with the OXPHOS complex V (ATP synthase) inhibitor, oligomycin, increased viral load and cytokine mRNA levels. Treatment with ROS scavengers N-acetyl cysteine (NAC) and MitoQ decreased HIF-1α protein levels, mRNA expression of pro-inflammatory cytokines, glycolytic and mitochondrial proteins, and viral load (*16*).

SARS-CoV-2 infected human peripheral blood mononuclear cells (PMBC) and CaCo-2 cells also increased transcription of the HIF-1α (*12*). SARS-CoV-2 infection of PBMCs showed down regulation of mtDNA transcripts, mitochondrial function, and increased glycolysis (*17*) in association with elevated mitokines, fibroblast growth factor 21 (FGF21)(*18*) and growth differentiation factor 15 (GDF-15)(*19*). Expression of FGF-21 and GDF-15 are increased when the mitochondrial unfolded protein response (UPR^MT^) and the cytosolic unfolded protein response (UPR^CT^) are activated causing an imbalance between the nDNA and mtDNA mitochondrial translation products (*20*).

Mitochondrial function is modulated by mTORC1 activity and mTORC1 is increased in COVID-19 lung tissue, upon SARS-CoV-2 infection of Vero cells, and in infected human bronchial epithelial cells. Activation of mTORC1 is known to increase glycolytic metabolism by activating c-Myc and stabilizing HIF-1α (*21, 22*), which impairs mitochondrial biogenesis (*22*). Conversely, the mTORC1 inhibitor rapamycin, which increases mitochondrial function, decreased viral replication in cell lines (*23*). Thus, SARS-CoV-2 infection impairs mitochondrial function, increases mROS production, and stabilizes HIF-1α which facilitates viral replication.

## SARS-CoV-2 Polypeptides Bind Mitochondrial Proteins

One mechanism by which SARS-CoV-2 manipulates mitochondrial function is by direct binding of viral polypeptides to host proteins. Up to 16% of host proteins bound to individual SARS-CoV-2 polypeptides in human HEK293T and A549 cells were mitochondrial (*13, 24, 25*) **(Table S1).** For example, the SARS-CoV-2 M protein interacted with the mitochondrial protein synthesis proteins leucyl-tRNA synthase (*TARS2*) and asparaginyl-tRNA synthase (*NARS2*); the coenzyme Q synthesis protein *COQ8B*; the N terminal imported protein peptidases PMPCA and PMPCB; and complex I subunits NDUFS2, NDUFB8 and NDUFB10. The SARS-CoV-2 non-structural protein, Nsp6, bound OXPHOS complex V (ATP synthase) subunits; SARS-CoV-2 open reading frame 10 (Orf10) bound the mitochondrial inner membrane protein import polypeptide, TIMM8; and Orf9b bound mitochondrial outer membrane protein import polypeptide, TOMM70 (*13, 24–27*) (**Table S1**).

While binding of viral polypeptides to host proteins could account for the mitochondrial effects of acute infection, SARS-CoV-2 infected K18-hACE2 mice showed broadly reduced nDNA and mtDNA encoded mitochondrial OXPHOS proteins in the heart, lung, kidney, and spleen (*28*). This implies a more global inhibition of mitochondrial biogenesis, which could occur at the transcriptional level throughout the infection cycle. To address this potentiality, we investigated the effects of SARS-CoV-2 infection on the transcript levels of mitochondrial OXPHOS, glycolysis, nutrient sensing, and stress response genes. We examined human early-stage infection nasopharyngeal swabs and late-stage infection autopsy samples of SARS-CoV-2 positive and negative individuals collected early in the pandemic in New York City (*29*). We then examined SARS-CoV-2 infected hamsters and mice to analyze changes in bioenergetic gene expression at early-stage and mid-stage infection.

These analyses revealed major changes in mitochondrial gene expression as the SARS-CoV-2 infection progressed. These changes proved to be due to two interacting factors, the concerted down regulation of groups of mitochondrial genes by the virus which is opposed by the host cell compensatory induction of mitochondrial gene expression. In the end, while the host compensatory response prevailed in the autopsy lung, it failed dramatically in the autopsy heart.

## RESULTS

We calculated the relative expression levels of host genes in RNA-seq data from ∼700 nasopharyngeal samples and ∼40 autopsy cases from SARS-CoV-2 positive and negative individuals, using the curated cellular bioenergetics genes listed in **Table S2** plus the gene and pathway lists from MitoCarta and MitoPathway (*30*). These results were validated and extended using hamsters and mouse models.

### Early-Stage SARS-CoV-2 Infection: Transcript Changes in Nasopharyngeal and Blood Samples

#### Mitochondrial Gene Expression Analysis of SARS-CoV-2 Infected Nasopharyngeal Samples

Analysis of the nasopharyngeal samples, revealed the interaction between viral suppression of mitochondrial gene expression and host compensatory mitochondrial gene induction (**Fig. 1**). Samples were binned into no, low, mediate, and high SARS-CoV-2 viral loads, and infections by other pathogens. The expression in the nDNA-coded genes for OXPHOS complexes I, II, III, IV, and V was significantly perturbed in the intermediate and high viral load samples exhibiting a striking dichotomy in responses-some OXPHOS genes being strongly down regulated (blue) and others strongly up regulated (red)(**Fig. 1A**).

**Fig. 1.**
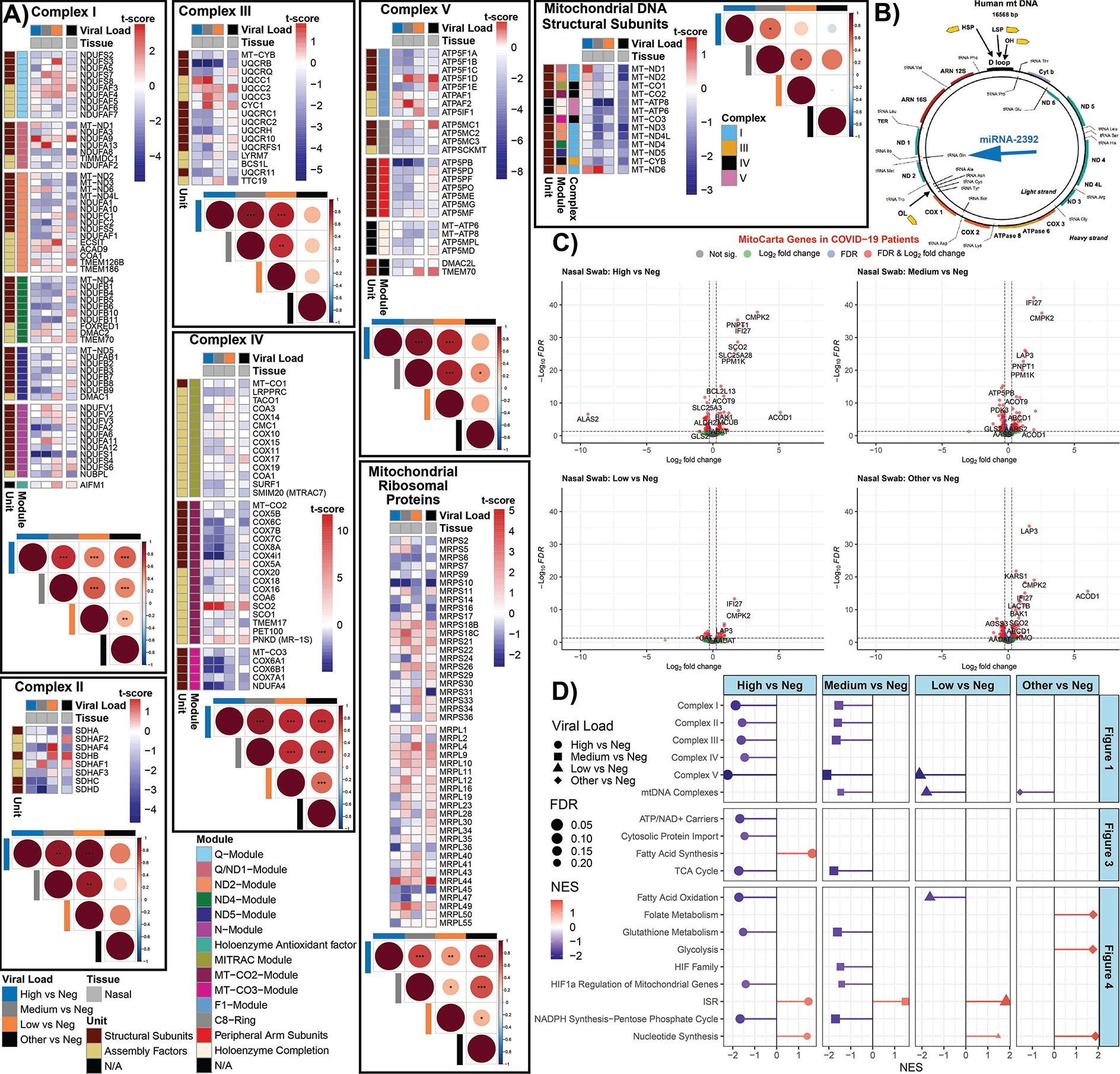
Mitochondrial OXPHOS complex module gene expression in nasopharyngeal samples from COVID-19 patients. **A)** Heatmaps of t-score statistics for specific mitochondrial complex genes expression. Correlation plots associated with each heatmap are shown for all the samples. Circle size is proportional to the correlation coefficient. Two-tailed Student’s t-test *p<0.05, **p<0.01, ***p<0.001. **B)** Schematic of the human mtDNA with the position that miR-2392 will interact. **C)** Volcano plots for differentially expressed genes related to mitochondrial processed (MitoCarta). **D)** Lollipop plots custom-curated mitochondria gene sets determined by fGSEA. NES, nominal enrichment score.

Each mitochondrial OXPHOS complex is composed of both nDNA and mtDNA coded proteins assembled in stoichiometric ratios via subassembly modules. Analysis of the differentially expressed genes indicated that SARS-CoV-2 infection coordinately suppressed the expression of genes associated within mitochondrial OPXPHOS complex assembly modules. This contrasted with the up regulation of mitochondrial genes not inhibited by the virus.

The mtDNA codes for 13 OXPHOS polypeptides plus the tRNAs and rRNAs necessary for the expression of these genes via mitochondrial protein synthesis. The remaining mitochondrially-localized proteins are nuclear coded, with the genes dispersed across the chromosomes. The nDNA transcripts are translated on cytosolic ribosomes and the polypeptides imported into the mitochondrion via transport through the outer mitochondrial membrane (TOMM) and inner mitochondrial membrane (TIMM) import complexes.

The multi-subunit OXPHOS complexes are assembled via intermediate stage modules, often requiring specific assembly factors (*31–33*). To obtain a better understanding as to how SARS-CoV-2 intervenes in mitochondrial bioenergetics, we analyzed the mRNA levels of the structural genes and assembly factors for each of the OXPHOS complex modules (**Table S2**).

Complex I is composed of six modules. Four modules nucleate around mtDNA coded polypeptides, MT-ND1, MT-ND2, MT-ND4, or MT-ND5. The remaining two modules, Q and N, contain only nDNA subunits (**Fig. 1A, Table S2**). The nDNA coded subunit genes incorporated into the mtDNA polypeptide nucleated modules revealed a striking result. Four of the five structural genes of the MT-ND2 module were down regulated by viral infection. In contrast, five of the six assembly factors were up regulated. All six of the structural subunit genes of the MT-ND4 module were down regulated, but assembly genes *FOXRED1* and *TMEM70* were up regulated. All of MT-ND5 module’s nDNA coded genes were down regulated, and seven of the eleven N module nDNA genes were down regulated. By contrast, two of the six nDNA genes of the MT-ND1 module were strongly up regulated and half of the nDNA genes of the Q module were up regulated and half down regulated.

Complex II subunits and assembly factors are all nDNA coded. Five of the eight were down regulated in the high viral load samples (**Fig. 1A**).

Complex III encompasses one mtDNA subunit, MT-CYB, and has a more linear assembly. Most complex III nDNA genes were down regulated except for the intermediate assembly steps encompassing *UQCC1, UQCC2, UQCC3*, and *CYC1* which were strongly up regulated.

Complex IV is assembled around mtDNA genes. All four nDNA genes of the MT-CO3 module were down regulated. A cluster of five genes of the MT-CO2 module were strongly down regulated, while assembly factor *SCO2* was strongly up regulated. MT-CO1 module gene expression was minimally affected.

Complex V transcript analysis revealed that all of the nDNA genes in modules ATP5MC1, ATP5PB, MT-ATP6/8, as well as the assembly factors *DMAC2L* and *TMEM70*, were down regulated, with the *ATP5PB* gene being especially suppressed. Expression of ATP5F1A module genes was variable.

Finally, the expression of the mitochondrial ribosomal genes was variable. The *MRPS10, MRPS16, MRPS34, MRPL36* genes were strongly down regulated while *MRPL44* was strongly up regulated (**Fig. 1A**). This parallel down regulation of multiple nDNA genes in individual OXPHOS modules suggests that these genes may be coordinately transcribed. If true, SARS-CoV-2 would only need to intervene in the common transcriptional factor(s) of one of the modules for each complex to disrupt OXPHOS biogenesis. In response, the host cell appears to counter by up regulating all the OXPHOS genes, however only those not blocked by the virus are induced.

This interaction between viral inhibition of the transcription of certain OXPHOS genes versus the host induction of all mitochondrial genes is corroborated by the unique expression profile of the mtDNA coded OXPHOS genes (**Fig. 1B**). In the samples with other viral infections or SARS-CoV-2 negative samples, all the mtDNA transcripts were proportionately down regulated. By contrast, in the samples with the highest SARS-CoV-2 RNAs, most mtDNA mRNAs were either unchanged or down regulated, with the exception of two mtDNA mRNAs, *MT-ND1* and *MT-ND6*, which were strongly up regulated. The mtDNA polypeptide genes are transcribed from two promoters as long polycistronic transcripts for each mtDNA strand, with the polypeptide genes punctuated by tRNA gene sequences. As the transcripts are synthesized, the tRNAs are cleaved out generating the mature mRNAs. For the G-rich “heavy” (H), one promoter transcribes 12 polypeptide genes. For the C-rich “light” (L) strand the other promoter transcribes only one polypeptide gene, *MT-ND6* (**Fig. 1B**).

In our companion paper (*34*), we reported that SARS-CoV-2 up regulates microRNA-2392 (miR-2392). It has been proposed that miR-2392 enters the mitochondrion and binds to the mtDNA tRNA *MT-TQ* gene sequence, impeding further progression of the mitochondrial RNA polymerase (*35*). We would thus anticipate that in COVID-19 the genes between the H strand promoter and *MT-TQ* would be induced while those downstream from *MT-TQ* to be reduced. This proves to be the case. The *MT-ND1* mRNA upstream of *MT-TQ* is strongly up-regulated relative to uninfected samples while the next three genes *MT-ND2, MT-CO1,* and *MT-CO2* have lower mRNA levels and the farther downstream polypeptide genes *MT-CO3* and *MT-CYB* were strongly down-regulated. The only other strongly up-regulated mtDNA gene is *MT-ND6* which is transcribed from the independent L-stand promoter.

To further clarify the effects of SARS-CoV-2 infection on mitochondrial function, we used volcano plots to broaden our analysis to the transcript levels of all MitoCarta identified mitochondrial proteins (**Fig. 1C**). In the high viral load samples, several mRNAs were strongly up regulated. Most striking was cytosine monophosphate kinase 2 (*CMPK2*). CMPK2 is the rate limiting step for mtDNA replication, being central to the creation of oxidized mtDNA (Ox-mtDNA) which is released from the mitochondrion and binds to the inflammasome (*36*). *PNPT1* (polyribonucleotide nucleotidyltransferase 1), also highly induced, is thought to be involved in processing mitochondrial double-stranded RNA released into the cytosol to engage MDA5-driven anti-viral signaling and trigger the type I interferon response (*37*). Also, up regulated was *IFI27* (Interferon Alpha Inducible Protein 27), which in blood is the transcript that provides the greatest discrimination between SARS-CoV-2 positive and negative patients (*38*), and *ACOD1*(Aconitate Decarboxylase 1 or *IRG1*), which regulates itaconate production and the metabolic adaptation of inflammatory myeloid cells (*39*). Additional up regulated genes included central energy metabolism genes including *SCO2* (cytochrome c oxidase, COX assembly), *SLC25A3* (phosphate carrier) critical for ATP synthesis, and genes for metabolism and for regulating apoptosis (**Table S3**).

Down-regulated genes include *ATP5PB* (ATP Synthase Peripheral Stalk-Membrane Subunit B); *PDK3* (Pyruvate Dehydrogenase Kinase 3); and additional metabolic genes (**Table S3**). Hence, SARS-CoV-2 infection shows a dose-response regulation of mitochondrial gene expression. In addition to the general inhibition of OXPHOS gene expression and mitochondrial metabolism, SARS-CoV-2 activates the mitochondrial innate immunity response (*CMPK2, IFI27* and *IRG1*) genes.

#### Pathway Analysis of Gene Expression Changes in SARS-CoV-2 Infected Nasopharyngeal Samples

To broaden our analysis of the transcriptional changes of cellular bioenergetic pathways during early nasopharyngeal infections, we performed Gene Set Enrichment Analysis (GSEA) on custom mitochondria pathways and MitoPathway created pathways (**Fig. 1D, Fig. S2**). At high viral loads, gene expression levels of the OXPHOS complexes I, II, III, and V were strongly down regulated along with mitochondrial cytosolic protein import complexes, mitochondrial TCA cycle and metabolic genes, fatty acid oxidation, glutathione synthesis, HIF-1α regulated mitochondrial genes, and the pentose phosphate shunt. By contrast, genes for fatty acid synthesis, nucleotide synthesis, and the integrated stress response (ISR) were up-regulated (**Fig. 1D**). Since pharmacological inhibition of fatty acid synthesis in SARS-CoV-2 infected hPSC-AO, HEK293T-hACE2 cells and K18-hACE2 mice inhibits viral replication (*14, 40*), induction of fatty acid biosynthesis must be required for viral replication, perhaps for the formation of the viral double membrane assembly structures (*41*).

The overall down regulation of OXPHOS complex IV genes is contrasted by the strong induction of the complex IV assembly gene, *SCO2*, (**Fig.1D**), consistent with a cellular compensatory effort to offset the viral reduction in complex IV gene transcription. The mitochondrial inner membrane nucleotide transport proteins (*SLC25A3* and *SLC25A4*) required for ATP synthesis were also down regulated, as were the mtDNA’s replication and transcription genes, other transmembrane solute carriers (*SLC25A1*), coenzyme Q (CoQ) synthesis genes which bind M, the inner and outer mitochondrial membrane polypeptide transport carrier genes including TOMM70 which binds Orf9b, and mitochondrial fission and fusion genes (**Fig. S2**).

By contrast, the hypoxia initiation factor regulatory genes were strongly up regulated, though on average the large number of HIF-1α target genes mitigates this effect. A number of the mitochondrial bioenergetic and biogenesis genes that were down-regulated at high viral loads were up-regulated at lower SARS-CoV-2 loads (**Fig. 1A**), perhaps again reflecting a cellular compensatory response to the early SARS-CoV-2 inhibition of mitochondrial function.

In summary, SARS-CoV-2 inhibits the transcription of a wide range of genes critical for the bioenergetic machinery, thus inhibiting cellular oxidation of substrates in favor of activation of HIF-1α. This likely directs substrates through the early stages of glycolysis to increase fatty acid synthesis to promote viral replication. In response, the cell up regulates transcription of nDNA and mtDNA bioenergetic genes as well as mitochondrial innate immunity genes (*CMPK2, IFI27* and *IRG1*) (*36, 42*).

#### Interaction Between SARS-CoV-2 Protein Inhibition and Transcript Inductions in Nasopharyngeal and Peripheral Blood Samples

To assess the potential functional effects of the changes in nasopharyngeal gene expression with functional protein changes, we compared changes in the acutely infected patient nasopharyngeal transcript levels with changes in the acutely SARS-CoV-2 infected human colon epithelial carcinoma cell line, CaCo-2 (*9*) (**Fig. S3A&B**). This revealed the strong transcriptional down regulation of the cytosolic ribosomal protein mRNAs, which would activate UPR^MT^, UPR^CT^, and ISR, though the ribosomal protein levels were not significantly changed in this short-term CaCo-1 infection experiment. Consistent with a marked down regulation of multiple OXPHOS mRNAs, CaCo-2 infected cells down regulated OXPHOS complex I (NADH dehydrogenase) in association with the strong increase in pyruvate dehydrogenase kinase, which negatively regulates pyruvate dehydrogenase, and proteins involved in increasing mitochondrial inner membrane permeability.

mtDNA transcripts and mitochondrial RNA polymerase (*POLRMT*) have been reported to be down regulated in blood cells while interferon response genes were induced (*17*) with the mitochondrial *IFI27* transcript being the most discriminatory factor in SARS-CoV-2 blood (*38, 43*). We observed in COVID19 positive cases, the blood mRNA levels for *FOXRED1* (complex I assembly) and *SCO2* (complex IV assembly) were up regulated (**Fig. S4A**) as we observed in complex I module MT-ND4 and in complex IV MT-CO2 module, *SCO2* induction also being observed by others (*17*) (**Fig. S4B**). The mRNA levels for the antioxidant enzyme, catalase (*CAT*) and for ferritin heavy chain (*FTH1*) for iron homeostasis, were reduced, though their protein levels in COVID19 positive case plasma were increased (*43*). The reduction in CAT transcription is consistent with the virus inhibiting OXPHOS and antioxidant enzymes to increase mROS levels (**Fig. S4B**)(*17*). Finally, mRNA and protein levels were up regulated for nDNA replication (*TYMS, TK1, PAICS*), folate metabolism (*DHFR*), and stress response (*DDIT4* regulation of mTOR).

### Late-Stage SARS-CoV-2 Infection and Lethality: Transcript Changes in COVID-19 Human Autopsy Samples

To determine the changes in bioenergetic gene expression late in infection, we analyzed the expression profile of patients who died of COVID-19. The autopsy samples from 35 deceased COVID-19 patients encompassing high and low SARS-CoV-2 levels were compared to five uninfected controls. Samples were collected from the heart, kidney, liver, lung, and lymph nodes (*44*). We also included publicly available RNA-seq data on monocytes from COVID-19 patients (*45*).

### Perturbation of OXPHOS Gene Expression in Autopsy Tissue of SARS-CoV-2 Infected Patients

Analysis of the transcriptional changes of mitochondrial OXPHOS bioenergetics and protein synthesis genes revealed a striking down regulation of virtually all OXPHOS genes in the heart (**Fig. 2A**). This included OXPHOS structural genes and assembly factors for complexes I, II, III, IV, and V as well as mitochondrial ribosomal proteins. This was not simply the product of terminal destruction of heart cells, since assembly genes for complex IV (*COX16, COX20, COA6, SCO2,* and PET100) were up regulated.

**Fig. 2.**
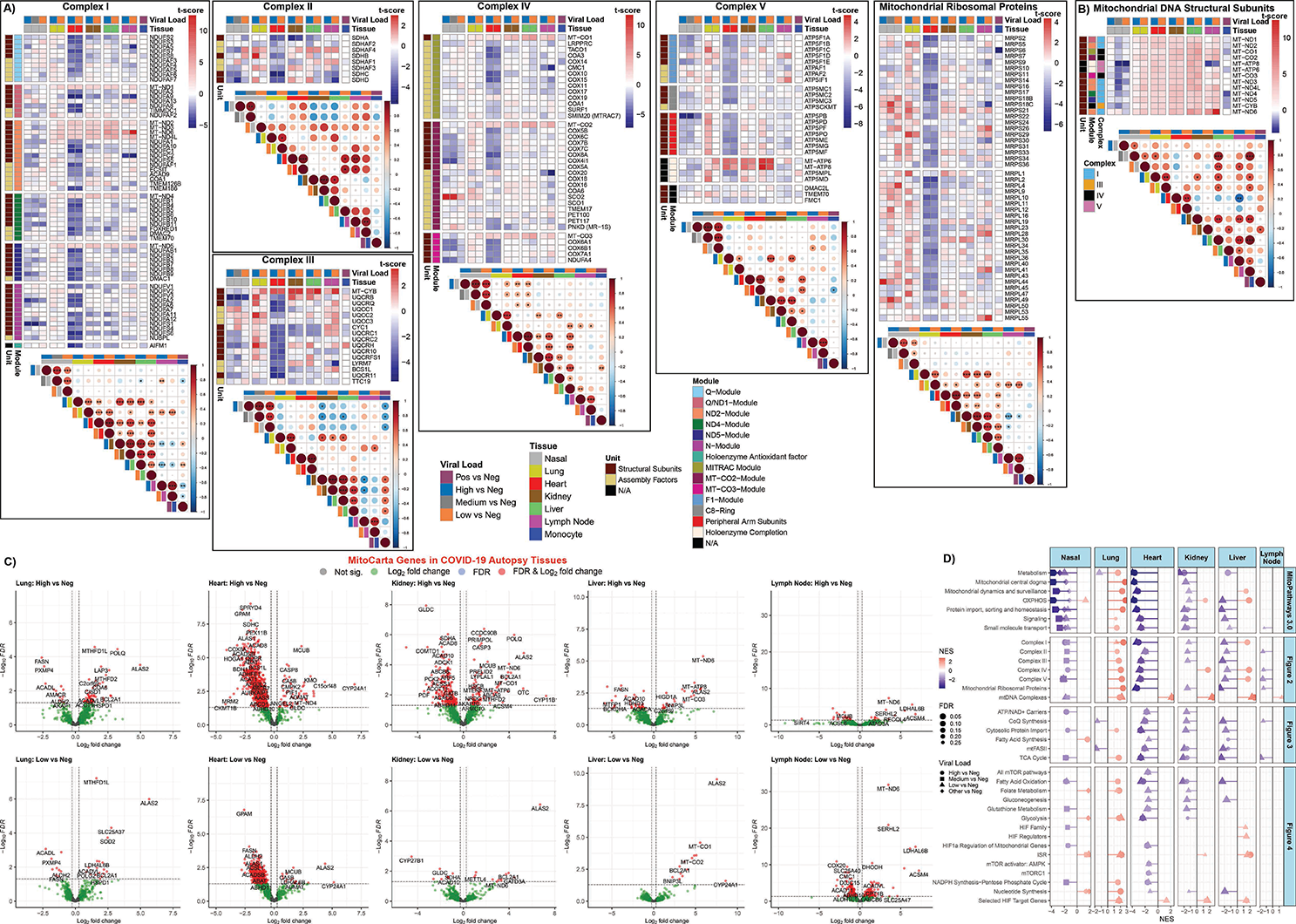
Mitochondrial OXPHOS complex module gene expression in COVID-19 patients from autopsy tissues. **A)** and **B)** Heatmaps display the t-score statistics when comparing viral load versus negative patient samples for nasopharyngeal swab and autopsy samples for specific mitochondrial complex genes. Correlation plots associated with each heatmap are shown for all the samples. Circle size is proportional to the correlation coefficient. Statistical significance determined using a two-tailed Student’s t-test *p<0.05, **p<0.01, ***p<0.001. **C)** Volcano plots for differentially expressed MitoCarta genes from autopsy COVID-19 patient samples. **D)** Lollipop plots for statistically significant custom mitochondria and MitoPathway gene sets determined by fGSEA. NES, nominal enrichment score.

The kidney and liver of SARS-CoV-2 infected patients showed down regulation of remarkably similar subsets of the OXPHOS genes, though many fewer genes than in the heart. These included the down regulation of selected complex I structure genes in the MT-ND2, MT-ND4 and NDUFV1 modules, complex II *SDHA*, and complex III structural and assembly genes *CYC1, UQCRC1, BCS1L and TTC19.* This further supports the possibility that SARS-CoVC-2 may interfere with specific transcriptional elements for OXPHOS genes.

Changes in the transcriptional profiles of the lymph nodes revealed individual complex genes being down regulated: *NDUFA5, NDUFA9, NDUFC2* in complex I modules NDUFS2, MT-ND1, and MT-ND2; *SDHC* and *SDHD* in complex II; and *CMC1* and *COX20* in complex IV modules MT-CO1 and MTCO2. Only modest changes were seen in OXPHOS gene expression in the monocytes (**Fig. 2A**).

Mitochondrial OXPHOS gene expression was almost uniformly up regulated in lung autopsy samples in striking contrast to other organs, as previously reported (*10*). Notably induced OXPHOS genes in lung included *SDHB* in complex II; *UQCRB, UQCRQ, UQCC2, UQCRH,* and *UQCRFS1* in complex III; *ATP5F1C* and all of the genes in the ATP5PB and the MT-ATP6/8 modules of complex V; as well as the majority of the genes for the mitochondrial ribosomal proteins (**Fig. 2A**). Possibly by the time the COVID-19 patients die from organ failure, the lungs have cleared the virus and the up regulation of OXPHOS gene expression comes to predominate. In contrast to the variable expression of the nDNA coded OXPHOS genes between tissues, all the mtDNA coded genes were strongly up regulated in all the autopsy tissues (**Fig. 2B**). This implies that miR-2392 is no longer inhibiting mtDNA transcription consistent with lower viral RNA levels, and confirms that the down regulation of OXPHOS genes in the heart, kidney and liver is not the result of generalized tissue and organ destruction.

The differential regulated mitochondrial genes in autopsy tissues is further exemplified by analysis of the MitoCarta genes relative to uninfected controls plotted in volcano plots (**Fig. 2C**). There was a clear progression in the extent of down regulated mitochondrial gene expression from most severe in the heart, followed by kidney and liver, and then lymph node. The lung showed the reverse bias with predominantly up regulation of mitochondrial genes (**Fig. 2C**). The reduced effects on mitochondrial gene expression between high and low viral loads of each tissue confirm that these gene expression changes are the result of viral infection.

To further elucidate the functional implications of the differential mitochondrial gene expression among the different tissues, we selected some of the most striking up and down regulated genes of each tissue and determined their functions (**Fig. 2C, Table S3**). In the COVID-19 heart the most strongly up regulate gene was *MUCB*, a component of the mitochondrial Ca^++^ uniporter, which coincides with induction of *CYP24A1* which regulates vitamin D and Ca^++^ and PO_4_^-^ levels. This implies that mitochondrial Ca^++^ regulation is an important aspect of SARS-CoV-2 pathogenesis. The rate limiting gene for mtDNA replication, *CMPK2*, was up regulated along with other mtDNA replication genes (*PIF-1* and *ANGEL2*). Additional up regulated heart genes included *C15orf48,* a COX protein that on viral infection displaces NDUFA4 plus generates miR-147b that inhibits *NDUFA4* translation and enhances *RIG-1/MDA-5* mediated viral immunity (*46*), as well as genes associated with apoptosis, mitochondrial metabolism, and mtDNA coded genes (**Fig. 2C, Table S3**).

The most strongly down regulated gene in the COVID-19 heart was *GPAM* (glycerol-3-phosphate acyltransferase), the commitment step of glycerolipid synthesis. An array of other mitochondrial OXPHOS, peroxisomal, and mitochondrial metabolism genes were also down regulated, including complex II subunit *SDHC*; complex III assembly factor *BCSIL*; and complex IV subunit *COX8A* (**Fig. 2C, Table S3**).

In the COVID-19 kidney, *MCUB* was up regulated as in heart. Specific mtDNA replication, transcription and repair genes; apoptosis genes, metabolic genes, regulatory genes, and mtDNA transcripts (*MT-ND6, MT-CO1, MT-ATP6*) were also up regulated (**Fig. 2C**).

The strongest COVID-19 down regulated gene in kidney was *GLDC* for glycine cleavage, a gene which is up regulated in the autopsy heart. Multiple mitochondrial genes were also down regulated including *SDHA*, a subunit of complex II parallel to the down regulation of SDHC in the heart; *ADCK1*, which regulates coenzyme Q biosynthesis; mitochondrial fatty acid synthesis; fatty acid oxidation genes; mitochondrial metabolic genes; and genes involved in mitochondrial biogenesis including *PDF*, mitochondrial polypeptide deformylase, and *GATB*, which converts the miss addition of glutamate on tRNA^Gln^ to glutamine (**Fig. 2C Table S3**).

In the COVID-19 autopsy liver, up regulated genes included the OXPHOS regulatory genes *C2orf69* and *HIGD1A* and mtDNA transcripts *MT-CO3, MT-ATP6,* and *MT-ND6*, as well as regulators of apoptosis and metabolism. Down regulated genes included cytosolic fatty acid synthesis (*FASN*), mitochondrial fatty acid synthesis, mitochondrial metabolism, and a mitochondrial fission protein (**Fig. 2C, Table S3)**.

For the COVID-19 autopsy lymph nodes, up regulated genes included mtDNA nucleoid genes (*ATAD3A* and *RECQL4*); fatty acid oxidation genes; and metabolism genes. Down regulated genes include the Ca^++^ uniporter (*MCUB*), which was the opposite of heart and kidney, and the Sirtuin that regulates glutamine metabolism (*SIRT4*) (**Fig. 2C, Table S2**).

For the COVID-19 autopsy lungs, OXPHOS genes *C2orf69* and *COA6* for complex IV assembly; single carbon metabolism genes *MTHFD1L* and *MTHFD2*; heme and iron-sulfur center enzymes; heat shock and apoptosis genes; peptidases; DNA repair genes; and fatty acid metabolism were up regulated. In contrast, fatty acid synthetase (*FASN*), fatty acid oxidation, peroxisomal, metabolic and mitochondrial regulatory genes were repressed (**Fig. 2C, Table S3**).

Hence, COVID-19 inhibited transcription of bioenergetic genes in the autopsy heart, kidney, liver, and lymph nodes. Selected genes participating in mitochondrial OXPHOS, fatty acid and amino acid metabolism, protein import, mitochondrial dynamics, and molecular transport were repressed several log-fold in the heart, as they were in the nasopharyngeal samples **(Fig. 2D, Fig. S2)**, and to a lesser extent progressively in the kidney, liver, and lymph node. By contrast, in the lung OXPHOS genes were coordinately induced.

This human COVID-19 gene expression data indicates that SARS-CoV-2 infection initially impairs mitochondrial gene expression in the nasopharyngeal tissues and then progresses to down regulation of mitochondrial gene expression in the heart, kidney, and liver leading to death. However, in the lung the virus appears to be cleared before death resulting in the surviving lung cells up regulating inducing mitochondrial gene expression.

#### Expression of OXPHOS-Associated Genes in COVID19

We next focused on transcriptional changes in processes that complement mitochondrial respiration: mitochondrial ATP/ADP-Pi exchange, TCA cycle, coenzyme Q biosynthesis, mitochondrial fatty acid synthesis (mtFASII), fatty acid oxidation, and the cytosolic protein import system (**Fig. 3**). As with the OXPHOS genes, the COVID-19 autopsy heart showed a systematic down regulation of transcripts for multiple mitochondrial functions, while the kidney and liver showed less robust inhibition of mitochondrial gene expression, which paralleled the down regulation of mitochondrial gene expression in the infected nasopharyngeal samples. Again, the COVID-19 autopsy lung showed the converse with the general induction of mitochondrial genes (**Fig. 3**).

**Fig. 3.**
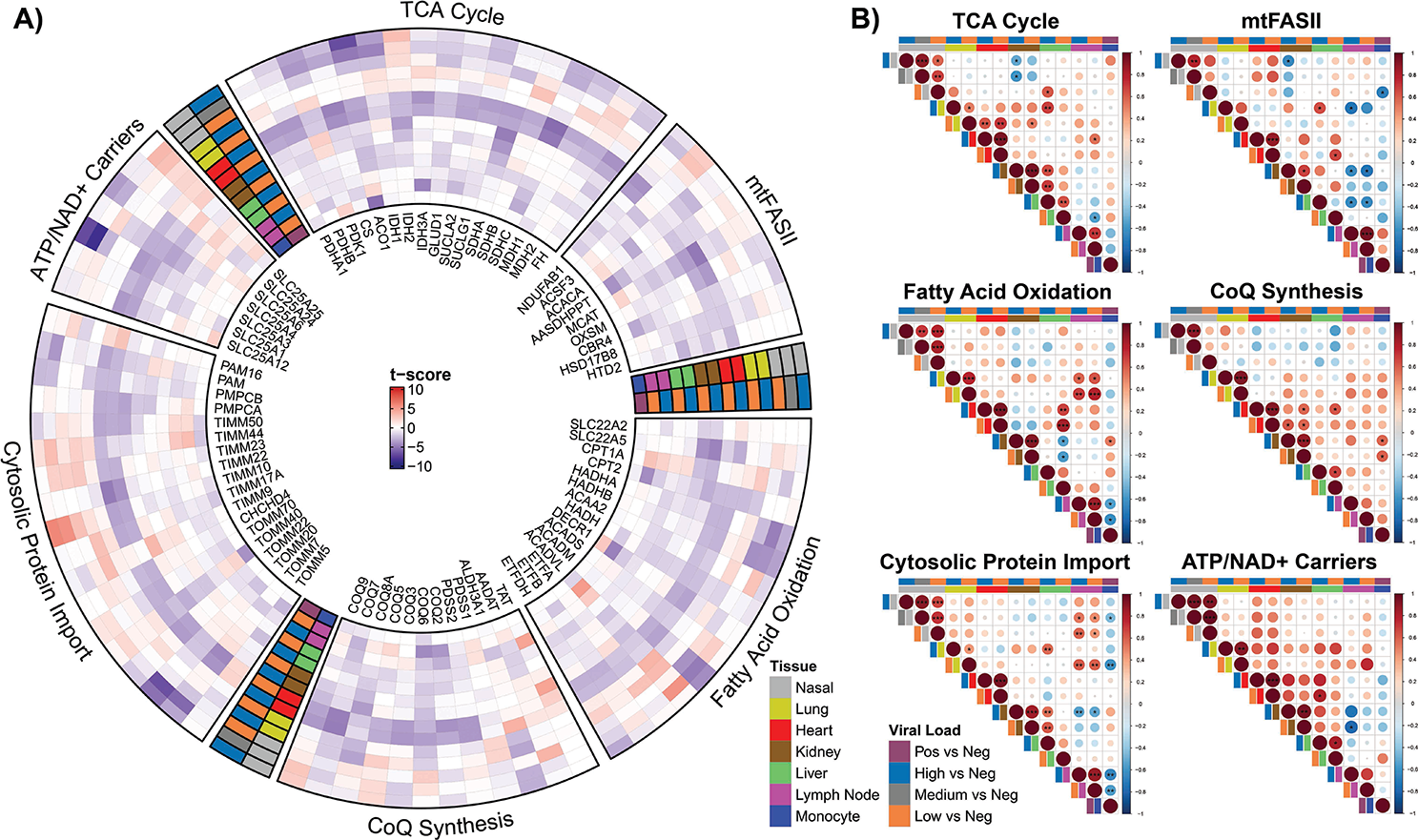
Analysis on mitochondrial ATP/ADP-Pi exchange, TCA cycle, coenzyme Q biosynthesis, mitochondrial fatty acid synthesis (mtFASII), fatty acid oxidation, and cytosolic protein import system. **A)** The circular heatmap displays the t-score statistics when comparing viral load versus negative patient samples for nasopharyngeal swab and autopsy samples for specific mitochondrial genes. **B)** Correlation plots associated with each group are shown for all the samples. Circle size is proportional to the correlation coefficient. Statistical significance determined using a two-tailed Student’s t-test *p<0.05, **p<0.01, ***p<0.001.

The differential expression of the mitochondrial inner membrane solute carrier genes is illustrative of the above dynamics. In the heart the phosphate carrier (*SLC25A3*), the adenine nucleotide translocases (ADP/ATP carries) (*SLC25A4*, ANT1) and *SLC25A6*, ANT3), the citrate carrier (*SLC25A1*), and the Ca^++^-binding aspartate-glutamate carrier (*SLC25A12*) were down regulated. By contrast, the ATP-Mg^++^/Pi transporters (*SLC25A24* and *SLC25A25*) were up regulated. *SLC25A3, SLC25A4, SLC25A6, SLC25A1, SLC25A12* are critical for mitochondrial energy production and ATP export, while *SLC25A24* and *SLC25A25* import ATP into the mitochondria without energetic benefit. The parallelism between the nasopharyngeal and heart samples is striking for these transport systems, with both tissues showing a pronounced down regulation of the phosphate carrier (*SLC25A3*) as well as ANT3 (*SLC25A6*) and up regulation of the ATP-Mg^++^/Pi transporter (*SLC25A25*). Thus, as the virus inhibits mitochondrial energetics, the cell responds by attempting to sustain intramitochondrial adenine nucleotide levels by importing ATP (**Fig. 3**).

#### Regulation of Mitochondrial Allied Bioenergetic Pathways

Changes in the expression of genes for fatty acid synthesis, NADPH synthesis, glutathione synthesis, folate metabolism, glycolysis, gluconeogenesis, and nucleotide metabolism exhibited a number of down regulated transcripts in heart and nasopharyngeal tissue, and generally the opposite in autopsy lung. However, the differences are much less pronounced than seen for the mitochondrial OXPHOS genes (**Fig. 4 A&B**).

**Fig. 4.**
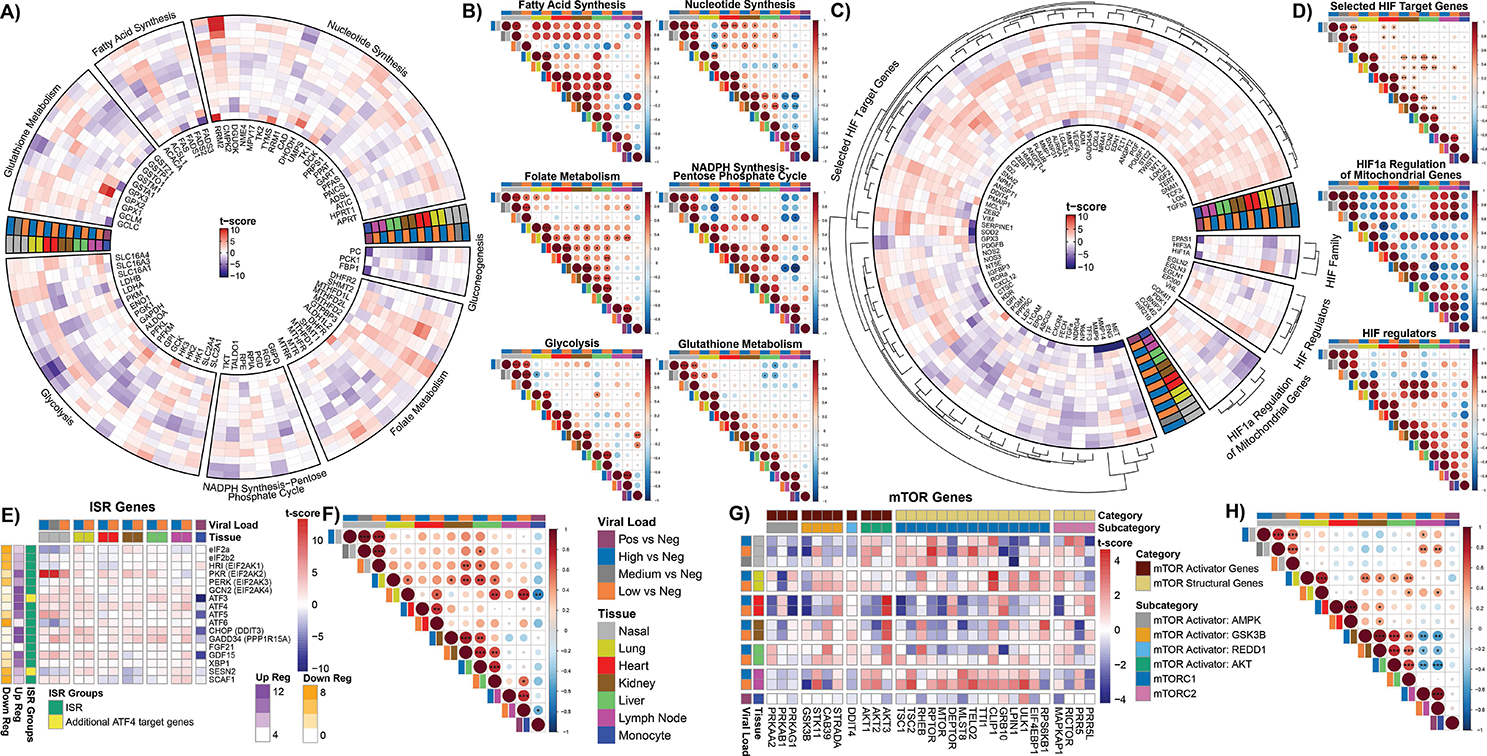
Glycolysis pathways, HIF pathways, ISR, and mTOR genes show differential expression. **A), C), E), and G)** Circular and tabular heatmaps display the t-score statistics when comparing viral load versus negative patient samples for nasopharyngeal swab and autopsy samples for genes related to specific pathways. The outer dendrogram in **C)** visualizes similar clusters as identified by complete-linkage clustering using Euclidean distance within each segment. **B), D), F), and H)** Correlation plots associated with each group are shown for all the samples. Circle size is proportional to the correlation coefficient. Statistical significance determined using a two-tailed Student’s t-test *p<0.05, **p<0.01, ***p<0.001.

Of particular interest was the expression of the single carbon metabolism genes which are metabolically interconnected between the mitochondrion and the nucleus-cytosol. We saw a general up regulation of all the mitochondrial folate enzyme genes, except dihydrofolate reductase 2 (*DHFR2*), in the heart, kidney, liver, lymph node verses the down regulation of the cytosolic and nuclear folate enzyme genes in the heart and progressively less so in kidney and liver. The lung pattern shows these trends but is more variable (**Fig. 4 A&B**). The up regulation of the mitochodnrial folate genes and the down regulation of the cytosolic and nuclear folate genes in the autopsy heart, kidney and liver (**Fig. 4A&B**) parallels the up regulation of mitochondrial single carbon metabolism seen in the cells and tissues of humans and mice with genetic defects in mitochondrial bioenergetics and biogenesis linked to activation of the UPR^MT^and the ISR (*47*).

In the nasopharyngeal tissue, we see the variable regulation of folate genes. However, we observed a striking induction of mtDNA replication gene, *CMPK2*, which has been linked to activation of the inflammasome (*36*).

For glycolysis in the heart, we observed the up regulation of lactate dehydrogenases A (*LDHA*) which converts pyruvate to lactate and favors glycolysis, but the down regulation of lactate dehydrogenase B (*LDHB*) which converts lactate to pyruvate and favors OXPHOS, a pattern also reported in peripheral blood (*17*). The glucose carriers (*SLC2A1* and *SLC2A4*) were down regulated as was the monocarboxylate carrier 1 (*SLC16A1*), which transports lactate across the mitochondrial inner membrane. However, monocarboxylate carrier 5 (*SLC16A4*) was strongly up regulated, as it is in cancer cells, being responsive to HIF-1α activation (*48*) (**Fig. 4 A&B**).

The down regulation of the glycolysis genes, *GAPDH* and *PGK1*, in the nasopharyngeal samples may seem anomalous, since we might have expected glycolysis to be induced to compensate for the mitochondrial ATP deficiency. However, inhibition of the later steps of glycolysis would drive carbon from the earlier steps of glycolysis into fueling fatty acid and NADPH synthesis which would generate the lipids critical for the production of the viral RNA replication centers.

In stark contrast to the down regulation of mitochondrial genes, the HIF-1α target genes were generally up regulated in autopsy tissues. This proved to be even more true for the autopsy lung in which HIF-1α was one of the most up regulated genes. This corresponds to the down regulation of *COX4i1* and the induction of *COX4i2* along with the mitochondrial LON protease which degrades *COX4i1*. This change shifts complex IV from a high oxygen to low oxygen optimal configuration (*49*). Perhaps, in the recovering lung cells, respiration is still impaired and maintaining the pseudohypoxic state is necessary to repair viral damage. This is again the opposite of the early infection nasopharyngeal tissues in which there is a variable expression of HIF-1α regulatory and target genes (**Fig. 4 C&D**).

#### Alterations in the ISR and mTOR Pathways in SARS-CoV-2 Infected Tissues

The differential inhibition of the cytosolic (**Fig. S3A**) and the mitochondrial (**Figs. 2 & 3**) ribosomes, plus increased mROS (*50*) would activate the UPR^MT^ and UPR^CT^ and the ISR via endoplasmic reticulum (ER) kinases HRI (EIF2AK1), PKR (EIF2AK2), PERK (EIF2AK3), and GCN2 (EIK2AK4)]. These kinases phosphorylate the cytosolic elongation factor, eIF2α, and phosphorylation of eIF2α impairs cytosolic protein synthesis and activates expression of the ISR transcription factors ATF4 and ATF5. Increased levels of ATF4 and ATF5 induce the expression of the downstream genes *GDF15*, *GADD34* and *CHOP* (*51, 52*). The ISR also induces the expression of the mitochondrial OXPHOS supercomplex assembly factor *SCAF1* (*53*) and the stress hormones FGF21 and GDF15 (*47*). While *GDF15* mRNA was up-regulated in nasopharyngeal, lung, and heart tissues, *FGF21* expression was not affected (**Fig. 4A)**.

*PKR*, which binds double standed RNA, was strongly up regulated in the nasopharyngeal samples along with *ATF3, ATF4, ATF5, CHOP, GADD34, GDF15,* while *eIF2α* was strongly down regulated. The autopsy heart, kidney, liver and lymph nodes showed a milder induction of the of the ISR genes, while the autopsy lung showed a mild up regulation of all of the ISR genes including *eIF2α*. Hence, during initial infection with high levels of SARS-CoV-2 replication intermediates, PKR is activated and induces the ISR. As the initial viral insult subsides, eIF2α is induced in lung, kidney, liver, and lymph node, presumably to return to normal cytosolic translation, though this does not happen in the heart (**Fig. 4E&F**).

SARS-CoV-2 infection is also associated with altered mTOR function, which also relates to mitochondrial dysfunction (*47, 53*). mTOR target genes were up regulated in the autopsy lung and lymph nodes, and to a lesser extent in the naropharyngeal samples (**Fig. 4G&H**). However, they were markedly down regulated in the autopsy heart, kidney, and liver. The induction of mTOR gene expression in the nasopharyngeal, lung, and lymph node tissues did not correlate with the induction of the AMPK genes (*PRKAA2, PRKAB1, PRKAG1*), but with the *STK11* kinase as well as *AKT1* and *AKT2.* By contrast, the down regulation of the mTOR genes in the heart, kidney, and liver correlated with the strong up regulation of *AKT3*. Presumably, in the heart, kidney, and liver, SARS-CoV-2 impairs mTOR gene expression which induces *AKT3*.

In summary, with the strong induction of *PKR* (*EIF2AK2*) in the acute infection nasopharyngeal samples down stream ISR genes were activated. ISR transcription was also activated in the other COVID-19 tissues. However, what distinguished the heart, kidney, and liver from the nasopharyngeal, lung and lymph node samples was the expression of the mTOR target genes which were down regulated in the former but up regulated in the latter. This corresponds to the differential induction of *STK11* plus *AKT1* and *AKT2* versus *AKT3* (**Fig. 4G&H**).

#### Utilization of Changes in COVID-19 Patient mRNA Levels to Investigate Viral-Induced Metabolic Fluxes

To further clarify the metabolic effects of SARS-CoV-2 acute nasopharyngeal and late-stage systemic infection we used the observed alterations in the host mRNA levels to estimate the flux rates of metabolites through bioenergetics via our computational model of cellular pathways (*54*). Analysis of mitochondrial bioenergetic and antioxidant defenses supported the conclusion that mitochondria function was down regulated in the nasopharyngeal samples and autopsy heart, kidney, liver and lymph node but strikingly up regulation in autopsy lung (**Fig. 5, Fig. S4**).

**Fig. 5.**
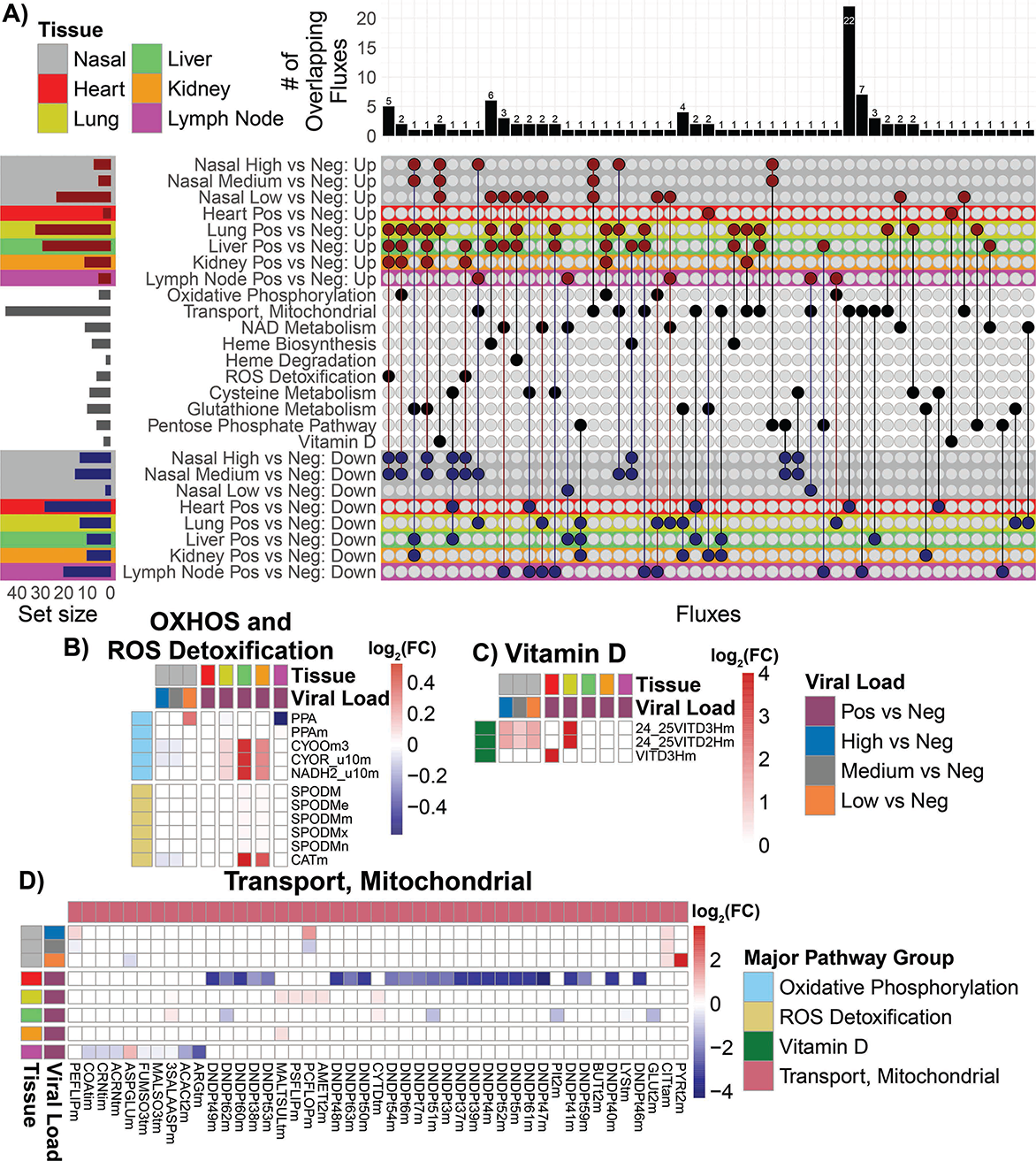
Metabolic flux analysis on RNA-seq data from nasopharyngeal and autopsy patient. **A)** Upset plot of the overlapping up regulated and down regulation metabolic fluxes with the nasopharyngeal and autopsy RNA-seq patient samples. The red color dots represent the up regulated metabolic fluxes and the blue color dots represent the down regulated fluxes. The bar chart on top are the number of overlapping metabolic fluxes for each intersect. The set size bar plot represents the total number of metabolic fluxes contained in each row. **B), C), and D)** Heatmaps of the of the log_2_ fold-change values for the significantly regulated fluxes in the OXPHOS, ROS detoxification, Vitamin D, and Mitochondrial Transport, major group flux pathways. The color bar represents the log_2_ fold-change values. The color of low but significant values are faint and may not be apparent in these heat maps.

In high and medium viral load nasopharyngeal samples, down regulation was observed for OXPHOS function; heme biosynthesis; ROS detoxification; and glutathione, cysteine, and pentose-phosphate metabolism. Similarly, autopsy heart samples showed a predominantly down regulation of mitochondrial function including 24 mitochondrial transport functions, far more than any other tissue, as well as reduced cysteine metabolism. Liver showed down regulation of six mitochondrial transport functions as well as glutathione and cysteine metabolism, while kidney exhibited down regulation five mitochondrial transport functions as well as glutathione and cysteine metabolism. Lymph nodes manifested down regulation of eight mitochondrial transporters and NAD, cysteine, and pentose phosphate metabolism.

The autopsy lung, by contrast, exhibited marked up regulation of OXPHOS; heme biogenesis and degradation; ROS detoxification; glutathione, cysteine and pentose-phosphate pathways; plus vitamin D metabolism. The liver and kidney also experienced increased heme biogenesis, ROS detoxification, glutathione metabolism, while liver exhibited increased heme metabolism, a few mitochondrial transport reactions, and cysteine and pentose phosphate pathways. The lymph nodes exhibited increased NAD metabolism and mitochondrial transport. For ROS detoxification, catalase was up regulated in liver and kidney, but down regulated in nasopharyngeal tissue. The only up regulated functions in the heart involved glutathione and vitamin D metabolism (**Fig. 5**).

Analysis of the modulation of the flux rates of the OXPHOS complexes is problematic due to their multiple subunit composition and the differential inhibition of the specific subunit transcripts by the virus versus induction of the remaining subunit genes by the host. Hence, high expression of some subunits by the host can overshadow the inhibition of others by the virus. Taking this into account, the heart had the lowest gene expression derived OXPHOS flux, while the kidney and liver, with increased calculated OXPHOS flux, had less severe viral inhibition of OXPHOS.

Consequently, calculated relative metabolic fluxes showed the same pattern as that observed for the individual mitochondrial genes. Nasopharyngeal and heart mitochondrial gene expression was strongly impaired; autopsy lung strongly induced; and liver, kidney and lymph nodes arrayed between.

### Early-Stage SARS-CoV-2 Infection in the Hamster Model: Analysis of Bioenergetic Gene Expression in Visceral and Brain Tissues

Analysis of the nasopharyngeal and autopsy samples revealed that mitochondrial gene expression is down regulated during early viral infection but is up regulated in later stages of infection when the virus is being cleared. To further elucidate the effects on mitochondrial function during early infection, we infected hamsters by nasopharyngeal administered SARS-CoV-2. When the virus reached its maximum titer in the hamster lung at four days post infection, we analyzed the mitochondrial gene transcript levels (**Fig. 6**). At this early time in infection, transcription of bioenergetics genes was not markedly perturbed in the lung, heart, kidney, and brain olfactory bulb. There were minimal changes in mitochondrial OXPHOS, mtFASII, coenzyme Q, protein import, TCA cycle, fatty acid oxidation, glutathione synthesis, and fatty acid synthesis gene expression. A subset of mitochondrial ribosomal genes were induced in the heart, lung and kidney, and mild increases were seen in heart and lung folate metabolism, NADP synthesis, nucleotide metabolism, HIF-1α target genes, and ISR genes.

**Fig. 6.**
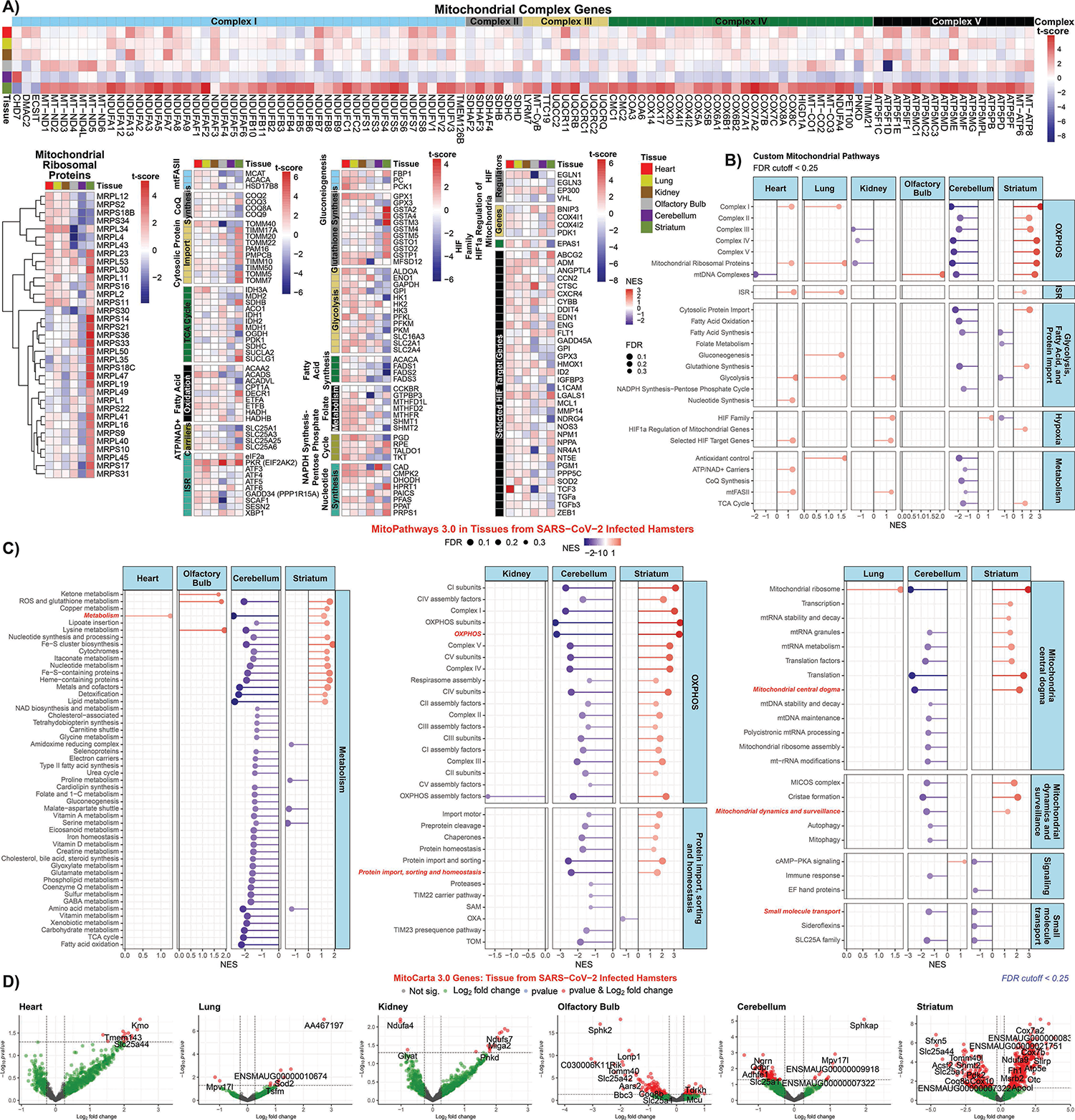
Mitochondrial specific analysis of RNA-seq data from SARS-CoV-2 infected hamster heart, lung, kidney, olfactory bulb, cerebellum, and striatum. **A)** Heatmaps display of mitochondrial specific genes with t-score statistics comparing SARS-CoV-2 infected hamsters vs. controls. **B) and C)** Lollipop plots for statistically significant custom mitochondria and MitoPathway gene sets determined by fGSEA. NES, nominal enrichment score. **D)** Volcano plots for differential MitoCarta genes from hamster tissues.

In stark contrast to the minimal effects of acute SARS-CoV-2 infection on the viscera, brain tissues showed striking changes in mitochondrial-related gene expression (**Fig. 6A,B,&C**). The cerebellum exhibited coordinate down regulation of bioenergetic processes including OXPHOS, mitochondrial ribosome, mtFASII, and cytosolic fatty acid synthesis and glycolytic genes. Conversely, the striatum showed a robust up regulation of OXPHOS, mitochondrial ribosomal, cytosolic protein import, glutathione synthesis genes, and a strong down regulation of glycolysis and cytosolic fatty acid synthesis genes

Volcano plot analysis of (**Fig. 6D**) revealed a bias toward increased mitochondrial gene expression in heart and kidney, presumably reflecting the high reliance of these organs on mitochondrial energy production; little perturbation of lung mitochondrial gene expression; general down regulation of olfactory bulb and cerebellum mitochondrial gene expression; and a marked increase in striatal mitochondrial gene expression. The functions of the genes highlighted in the volcano plots are listed in **Table S3**. Notable up regulated genes in the heart and kidney include the branched chain amino acid transporter (*Slc25a44*) in the heart and the complex I subunit (*Ndufs7*) and mitochondrial fusion protein (*Miga2*) in the kidney. Notable down regulated genes in the lung included the mitochondrial translational factor gene (*Tsfm*) and the mitochondrial superoxide dismutase (*Sod2*) gene.

The mitochondrial Ca^++^ uniporter gene (*Mcu*) was up regulated in the olfactory bulb, which parallels the induction of the mitochondrial Ca^++^ uniporter gene *MCUB* in the autopsy heart and kidney. Down regulated mitochondrial genes included the mitochondrial ATP synthase subunit 5 (*Atp5p*), mitochondrial alanyl-tRNA synthetase (*Aars2*), and citrate carrier (*Slc25a1*). In the cerebellum, the mitochondrial antioxidant gene *Mpv171* was up regulated while several neurological and mitochondrial genes were down regulated including the citrate carrier (*Slc25a1*). Finally, in the striatum, an array of mitochondrial genes was up regulated including complex IV subunit (*Cox7a2*), complex I subunits (*Ndufa9* and *Ndufb6*), TCA cycle fumarate hydratase-1 (*Fh1*), and mtDNA complex I subunit (*MT-ND3*). Notable striatal down regulated genes included citrate carrier (*Slc125a1*), mitochondrial folate serine hydroxymethyltransferase 2 (*Shmt2*), coenzyme Q synthesis (*Co8b*), and lactate dehydrogenase D (*Ldhd*).

These hamster results indicate that early in viral infection, when SARS-CoV-2 gene expression in the lung peaks and the viral polypeptides are binding to mitochondrial proteins to disrupt mitochondrial OXPHOS, the virus has yet to strongly influence host lung or visceral mitochondrial gene transcription. Surprisingly, however, we observed a striking differential expression of the brain bioenergetic genes, presumably responding to diffusible factors elaborated in response to viral infection, possibly impacting neurological function.

### Mid-Stage SARS-CoV-2 Infection in BALB/c and C57BL/6 Mouse Models: Analysis of Bioenergetic Gene Expression in the Lungs

To investigate the effects on mitochondrial function during a more advance stage of SARS-CoV-2 infection in the lung, we analyzed the mitochondrial transcripts of the lungs of BALB/c and C57BL/6 mice infected with mouse-adapted SARS-CoV-2MA10 virus at four days post-infection (*55*). BALB/c mice are more susceptible to lung pathologies and dysfunction caused by SARS-CoV-2 MA10 compared to C57BL/6 mice (*55*), and examining tissue in this time point provided data on the lung’s induction of mitochondrial gene expression as viral load declines.

We observed marked differences in the up and down regulated MitoCarta genes when comparing the two infected mouse strains (**Fig. 7A&B**). We found a more robust induction of OXPHOS complex I, II, III, IV, and V genes in the BALB/c compared to the C57BL/6 lungs. Similar patterns emerged for genes involved in mitochondrial protein import, TCA cycle, fatty acid oxidation, and inner membrane transport (including the ADP/ATP translocases), mitochondrial ribosomal genes, and ISR genes. Up regulation of genes involved in glycolysis, glutathione synthesis, folate metabolism, pentose-phosphate shunt, nucleotide synthesis, and HIF-1α target genes which were more intense in the BALB/c mice than the C57BL/6 mice. Finally, the cytosolic fatty acid synthesis genes, particularly genes that encode fatty acid synthetase (*Fasn*), acetyl CoA-carboxylase alpha (*Acaca*), and long-chain fatty acid ligase 1 (*Acsl1*), were more down regulated in the BALB/c than the C57BL/6 mice (**Fig. 7B**).

**Fig. 7.**
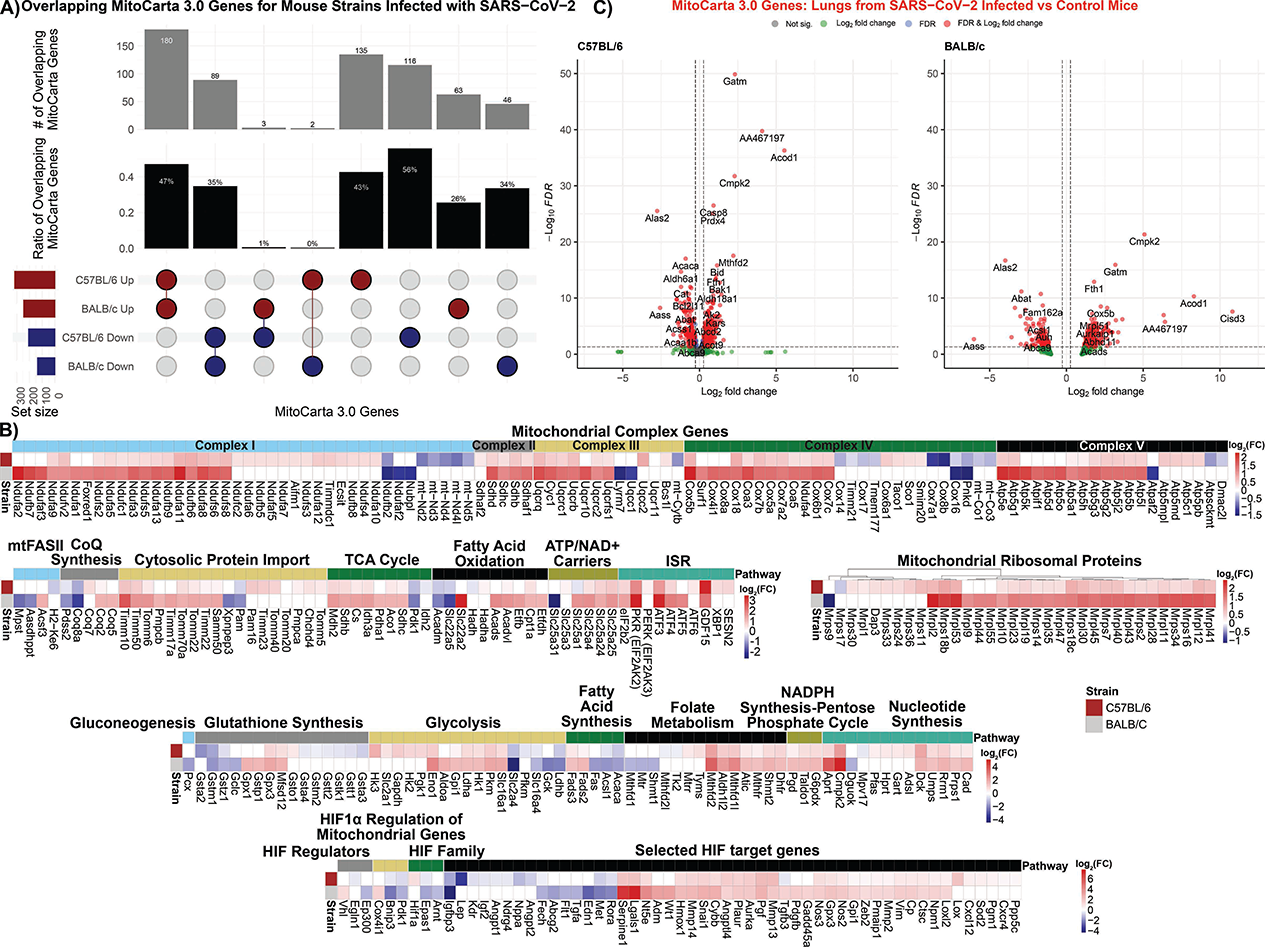
Mitochondrial specific analysis on RNA-seq data from SARS-CoV-2 infected mouse lungs. **A)** Upset plot of the overlapping MitoCarta genes comparing lungs from BALB/c and C57BL/6 mice infected with SARS-CoV-2. **B)** Heatmaps display of mitochondrial specific genes with t-score statistics comparing SARS-CoV-2 infected mice vs. controls. **C)** Volcano plots for differential MitoCarta genes from C57BL/6 and BALB/c tissues.

Volcano plots (**Fig. 7C**) and gene-level analysis of the mouse expression data revealed similar expression changes to the human autopsy lung. Among the top three to four induced genes of both mouse strains were *Cmpk2* and *Acod1*, involved in the mitochondrial modulation of inflammation, and *Gatm* which is required for creatin synthesis. Both strains experienced an up regulation of a range of mitochondrial metabolic and biogenesis genes, but the down regulation of *Alas2* involved in porphyrin synthesis (**Fig. 7C, Table S3)**.

Assuming that the C57BL/6 and BALB/c mice represent different stages of recovery from the viral infection of the lung, the C57BL/6 mice may be caught in the transition as the viral effects decline and the host induction is initiated. The BALB/c mouse may represent further disease progression with greater viral reduction and host gene induction, analogous to the human autopsy lung tissues.

## DISCUSSION

While previous studies have demonstrated that SARS-CoV-2 can inhibit mitochondrial function by the binding of viral structural, Nsp, and Orf proteins to critical host mitochondrial proteins (*13, 24, 25*) our current results demonstrate that a much more global mitochondrial inhibitory effect occurs at the transcriptional level. Nasopharyngeal and autopsy heart, kidney, and liver samples with high SARS-CoV-2 RNA levels manifest inhibition of transcription of mitochondrial genes associated with OXPHOS complexes I, II, III, IV, and V; fatty acid oxidation; antioxidant defenses; mitochondrial translational machinery; cytosolic protein import; mtFASII; mtDNA biogenesis; and intermediate metabolism. This contrasts with the autopsy lung which shows the opposite transcriptional profile, a general up regulation of mitochondrial gene expression. This striking difference demonstrates that the mitochondrial gene expression profiles changed during the course of the viral infection.

The relationship between viral load inhibition of mitochondrial transcriptional versus host compensatory induction is demonstrated by the differential regulation of nDNA OXPHOS genes in the nasopharyngeal tissues. The OXPHOS enzyme complexes are assembled from multiple nDNA and mtDNA coded subunits assembled through sub-enzyme modules which are then combined into the final enzyme. To assemble the polypeptides for each nodule in stoichiometric ratio, the modular genes would need to be coordinately regulated. In such a model, SARS-CoV-2 would need only to compromise the master transcription regulatory factor(s) for one enzyme complex module to block assembly of the enzyme. The transcriptional profiles of the OXPHOS modules in the SARS-CoV-2 infected nasopharyngeal samples supports the hypothesis of the coordinate regulation of the OXPHOS module genes and that SARS-CoV-2 blocks modular assembly by impacting the master transcriptional regulator of the modules nDNA genes. The host cells counter the viral inhibition by the coordinated up regulation of nDNA mitochondrial gene expression, demonstrated by the striking up regulation of the complex IV assembly gene, SCO2. Other pathway genes show potential coordinate modulation, such as the down regulation of the cytosolic-nuclear folate genes in association with the up regulation of the mitochondria folate genes.

SARS-CoV-2 manipulation of transcriptional pathways to impair mitochondrial bioenergetics is exemplified by the viral regulation of expression of the nasopharyngeal mtDNA transcripts. The SARS-CoV-2 genome codes for three sequences that are homologous to the seed sequences of miR-2392 (*34*). If these generate RNAs that mimic miR-2392, then at high SARS-CoV-2 viral loads, there would be increased inhibition of the genes and RNAs that bind to the miR-2392 seed sequence. It was previously proposed that miR-2392 binds to the mtDNA *MT-TQ* gene resulting in inhibition of mtDNA transcription. At high SARS-CoV-2 RNA levels, sufficient miR-2392 mimic RNA would be generated to enter the mitochondrion and block mtDNA transcription. This conjecture was supported by the high expression of the mtDNA H strand *MT-ND1* gene upstream from *MT-TQ*, and the reduced expression of the downstream mtDNA H strand mRNAs. SARS-CoV-2 manipulation of regulatory genes was further supported by the altered gene expression of the mTOR nutrient sensing pathway genes in association with the energy sensing kinases.

In the nasopharyngeal and heart, kidney, and liver samples, inhibition of OXPHOS and limitation of antioxidant defenses would result in increased mROS which stabilizes HIF-1α. This redirects metabolites away from the mitochondrion and toward glycolysis to generate lipid precursors. The imbalance in nDNA and mtDNA mitochondrial polypeptides also activates the UPR^MT^ and UPR^ER^, which activates the ISR resulting in bias of protein synthesis away from cellular maintenance and toward vial biogenesis.

The human autopsy data confirms that, as viral titer declines, normal mitochondrial function resurges repairing tissue damage. However, if the virally-induced inhibition has been too severe, as in the case of the autopsy heart, kidney, and liver, then organ failure ensues resulting in death.

While the early and late phases of COVID-19 modulation of bioenergetic gene transcription were established from the human nasopharyngeal and autopsy studies, the relationship between the initial viral protein inhibition of host mitochondrial proteins and the viral inhibition of bioenergetic gene transcription was not clear. To address this discrepancy, we analyzed the transcriptional effects of early hamster nasopharyngeal infection. This revealed that at peak lung viral titer, mitochondrial gene expression was not markedly impaired in the lung, heart, and kidney. Hence, the viral regulation of host mitochondrial transcription occurs after the initial viral protein inhibition of mitochondrial protein function. Surprisingly, however, brain mitochondrial gene expression was affected possibly accounting for the commonly experienced “brain fog” (*1*).

Late in the course of SARS-CoV-2 lung infection, an upsurge of bioenergetic gene expression occurred in the autopsy lung, likely associated with the clearance of the virus from the autopsy lung. To further clarify the host cell response to the progressive lung clearance of the virus, we studied the mid phase response of the lungs of C57BL/6 and BALB/c mice, C57BL/6 being more resistant than BALB/c mice to SARS-CoV-2MA10. The BALB/c lung showed the strong up regulation of mitochondrial gene expression seen in the human autopsy lung. However, the C57BL/6 mice appeared to be at an earlier disease stage, with the OXPHOS genes beginning to be induced, but mtDNA genes, OXPHOS assembly genes, and fatty acid oxidation genes still being down regulated.

Therefore, the most immediate bioenergetic effects of SARS-CoV-2 infection occur through the binding of viral polypeptides to critical host cell mitochondrial proteins disrupting mitochondrial metabolism and energetics. Next, the virus initiates a more global attack on mitochondrial bioenergetics by inhibiting the transcription of critical OXPHOS genes, a broad array of other mitochondrial genes, but at the same time preserving functions such as mitochondrial folate metabolism important for viral biogenesis. The global inhibition of mitochondrial OXPHOS would increase mROS which stabilizes HIF-1α resulting in the redirection of substrates away from mitochondrial oxidation and towards glycolysis to generate viral precursors. The host responds by the coordinate induction of bioenergetic gene expression but is foiled by viral blockage at critical transcriptional control nodes. The host also up regulates the mitochondrial innate immune genes (*CMPK2*, *IFI27*, *ACOD1/IRG1*), which contributes to the ultimate elimination of the virus, but exacerbating the host inflammatory response.

The host also responds by the coordinate induction of bioenergetic gene expression in other tissues but is foiled by viral blockage at critical transcriptional control nodes. While in the final stages of infection the lung succeeds in fully inducing mitochondrial transcription, this fails in the heart resulting in the comprehensive suppression of virtually all cardiac mitochondrial gene expression. This may explain why cardiac dysfunction is a prominent finding in COVID-19 (*2–5*). Since the heart is highly reliant on mitochondrial bioenergetics, protracted inhibition of mitochondrial bioenergetics could be an important factor on COVID-19 associated deaths.

The pivotal role of the mitochondrion in these processes suggests that an effective approach to mitigating the adverse effects of SARS-CoV-2 might be stimulation of mitochondrial function in combination with inhibition of mROS production. The beneficial effects of reducing mROS have been shown by the reduction of the HIF-1α protein levels, pro-inflammatory mRNA levels, and viral load by treating SARS-CoV-2 infected monocytes with the antioxidants N-acetyl-cysteine and MitoQ (*16*). The potential beneficial effects of up regulating mitochondrial function have been suggested by the rapamycin inhibition of mTORC1 which decreased viral replication (*23*). Viral pathology might also be mitigated by neutralizing miR-2393 (*34*) and activation of mitochondrial biogenesis with bezafibrate (*56*).

### Limitations of the Study

This study has a number of limitations. First, some of the clinical components of the study were designed retrospectively. Consequently, COVID-19 and control groups are not matched for demographic variables, comorbidities, or in-hospital treatments.

## Acknowledgments

This work was supported by supplemental funds for COVID-19 research from Translational Research Institute of Space Health through NASA Cooperative Agreement NNX16AO69A (T-0404 to A.B.) and further funding from KBR, Inc provided to A.B. This work used resources services, and support provided via the COVID-19 HPC Consortium (https://covid19-hpc-consortium.org/), provided specifically by the NASA High-End Computing (HEC) Program through the NASA Advanced Supercomputing (NAS) Division at Ames Research Center which was awarded to A.B.

Additional support came from DOD grant W81XWH-21-1-0128 awarded to D.C.W.

## Author contributions

Conceptualization: DCW and AB

Methodology: AB

Literature and concept integration: JWG, JMD, DCW, YSA, DGM, AA, LNS, SW, SMB, MRE

Formal Analysis: AB, HF, DT, MSK, HC, VZ, CM, CM, JCS, US, AS, JF, YZ, YHK, YZ

Investigation: AB, DCW, CEM, RES

Sample Collection: RES, CEM, MTH, NJM, EAM, SATB, EJA, WAS, RJD, BRT, JF

Resources: AB, CEM, RS, JCS

Writing – Original Draft: DCW, JWG

Writing – Review & Editing: AB, JWG, YSA, WP, SBB, RM, ESW, DT, RES, JCS, CEM, PMMV, VZ, MRE, MSK, YHK

Visualization: AB, HF, CM, MSK, HC, JWG, VZ Supervision: DCW and AB

Funding Acquisition: DCW and AB

## Data and material availability

The published article includes all datasets generated and analyzed during this study. Processed bulk RNA-seq data for the human related data from the nasopharyngeal and autopsy data is available online with dbGaP Study Accession number: phs002258.v1.p1 and online here at: https://www.ncbi.nlm.nih.gov/projects/gap/cgi-bin/study.cgi?study_id=phs002258.v1.p1 and also https://covidgenes.weill.cornell.edu/. The hamster and murine RNA-seq data is deposited on SRA (https://www.ncbi.nim.gov/sra), IDs pending. This study did not generate new unique reagents.

## Supplementary Materials for

### Materials and Methods

#### EXPERIMENTAL MODEL AND SUBJECT DETAILS

##### Human nasopharyngeal swab sample collection for RNA-seq analysis

Patient specimens were processed as described in Butler et al.(*1*). Briefly, nasopharyngeal swabs were collected using the BD Universal Viral Transport Media system (Becton, Dickinson and Company, Franklin Lakes, NJ) from symptomatic patients. Total Nucleic Acid (TNA) was extracted using automated nucleic acid extraction on the QIAsymphony and the DSP Virus/Pathogen Mini Kit (Qiagen).

##### Human autopsy tissue collection for RNA-seq analysis (5-8 controls & 35-36 patients)

The full methods of the patient sample collection from the autopsy patients are currently available in the Park et al. (*2*). All autopsies are performed with the consent of the next of kin and permission for retention and research use of tissue. Autopsies were performed in a negative pressure room with protective equipment including N-95 masks; brain and bone were not obtained for safety reasons. All fresh tissues were procured prior to fixation and directly into Trizol for downstream RNA extraction. Tissues were collected from lung, liver, lymph nodes, kidney, and heart as consent permitted. For GeoMx, RNAscope, trichrome and histology tissue sections were fixed in 10% neutral buffered formalin for 48 hours before processing and sectioning. These cases had a post-mortem interval of less than 48 hours. For bulk RNA-seq tissues, post-mortem intervals ranged from less than 24 hours to 72 hours (with 2 exceptions - one at 4 and one at 7 days - but passing RNA quality metrics) with an average of 2.5 days. All deceased patient remains were refrigerated at 4°C prior to autopsy performance. Host profiling was done on 39 patients that died from COVID-19 and healthy controls from organ donor remnant tissues (n = 3).

##### Virus lines

SARS-CoV-2 Washington strain (isolate USA-WA1/2020, NR-52281) were provided by the Center for Disease Control and Prevention and obtained through BEI Resources, NIAID, NIH. Cell lines Vero E6 (African green monkey [Chlorocebus aethiops] kidney, CVCL_0574; female) were obtained from ATCC (https://www.atcc.org/). Cells were cultured in Dulbecco’s Modified Eagle Medium (DMEM) supplemented with 10% FBS and 100 U/mL penicillin and 100mg/mL streptomycin. Cells were tested for the presence mycoplasma bi-weekly using MycoAlert Mycoplasma Detection Kit (lonza), They were not authenticated by an external service but were derived directly from ATCC.

##### SARS-CoV-2 propagation and infections

SARS-CoV-2 was propagated in Vero E6 cells in DMEM supplemented with 2% FBS, 4.5 g/L D-glucose, 4 mM L-glutamine, 10 mM Non-Essential Amino Acids, 1 mM Sodium Pyruvate and 10 mM HEPES using a passage-2 stock of virus. Three days after infection supernatant containing propagated virus was filtered through an Amicon Ultra 15 (100 kDa) centrifugal filter (Millipore Sigma) at 4000 rpm for 20 minutes. Flow through was discarded and virus was resuspended in DMEM supplemented as described above. Infectious titers of SARS-CoV-2 were determined by plaque assay in Vero E6 cells in Minimum Essential Media supplemented with2% FBS, 4 mM L-glutamine, 0.2% BSA, 10 mM HEPES and 0.12% NaHCO3 and 0.7% agar. All work involving live SARS-CoV-2 was performed in the CDC/USDA-approved BSL-3 facility with institutional biosafety requirements.

##### SARS-CoV-2 hamster model

Three-5-week-old male Golden Syrian hamsters (Mesocricetus auratus) were obtained from Charles River. Hamsters were acclimated to the CDC/USDA-approved BSL-3 facility for 2-4 days before viral infection. The hamsters were housed in a temperature-controlled room (20-22°C), on a 12 h light/dark cycle (08:00 – 20:00 lights on), with food (PicoLab Rodent Diet 20 5053) and water provided ad libitum. All hamsters were in good health and demonstrated normal behavior until the infection. All animal experiment procedures, breeding and ethical use were performed in accordance with the guidelines set by the Institutional Animal Care and Use Committee. Hamsters were anesthetized by intraperitoneal injection with a ketamine HCl/xylazine solution (4:1) before being intranasally inoculated with 100 pfu of SARS-CoV-2 isolate USA-WA1/2020 in PBS (or PBS only as a control) in a total volume of 100 ml. All animal experiments were performed on at least two separate occasions. Blinded treatment groups used mice throughout the study limited investigator subjectivity.

##### SARS-CoV-2MA10- murine model

All experiments were conducted using approved standard operating procedures and safety conditions for SARS-CoV-2 in BSL3 facilities designed following the safety requirements outlined in the Biosafety in Microbiological and Biomedical Laboratories, 6th edition, the United States Department of Health and Human Services, the Public Health Service, the Centers for Disease Control and Prevention (CDC), and the National Institutes of Health (NIH). All experiments and researchers followed protocols approved by the Institutional Animal Care and Use Committee (IACUC) at The University of North Carolina (UNC), Chapel Hill, NC.

The mice were housed in the UNC ABSL3 facility on a 12:12 light cycle using autoclaved cages (Tecniplast, EM500), irradiated Bed-o-Cob (ScottPharma, Bed-o-Cob 4RB), ad libitum irradiated chow (LabDiet, PicoLab Select Rodent 50 IF/6F 5V5R), and autoclaved water bottles. Animals used in this study included female 16-week-old C57BL/6J (B6) (The Jackson Laboratory stock 000664) or 10-12-week-old BALB/cAnNHsd (BALB/c) (Envigo order code 047) mice, purchased directly from vendors.

Mice were infected following light sedation, using 50 mg/kg ketamine and 15 mg/kg xylazine, by intranasal inoculation with 10^4^ pfu SARS-CoV-2-MA10 (*3*) diluted in 50 μL PBS or PBS alone (mock infection). Blinded treatment groups used mice throughout the study limited investigator subjectivity. Mice were euthanized by an overdose of isoflurane anesthetic and lung tissues collected for subsequent processing.

#### METHOD DETAILS

##### RNA-seq of nasopharyngeal swab COVID-19 patient samples

RNA isolation and library preparation are fully described in Butler, et al. (*1*). Briefly, library preparation on all the nasopharyngeal swab samples’ total nucleic acid (TNA) were treated with DNAse 1 (Zymo Research, Catalog # E1010). Post-DNAse digested samples were then put into the NEBNext rRNA depletion v2 (Human/Mouse/Rat), Ultra II Directional RNA (10 ng), and Unique Dual Index Primer Pairs were used following the vendor protocols from New England Biolabs. Kits were supplied from a single manufacturer lot. Completed libraries were quantified by Qubit or equivalent and run on a Bioanalyzer or equivalent for size determination. Libraries were pooled and sent to the WCM Genomics Core or HudsonAlpha for final quantification by Qubit fluorometer (ThermoFisher Scientific), TapeStation 2200 (Agilent), and qRT-PCR using the Kapa Biosystems Illumina library quantification kit.

##### RNA-seq of COVID-19 autopsy tissue samples

RNA isolation and library preparation is detailed in Park, et al.(*2*). Briefly, autopsy tissues were collected from lung, liver, lymph nodes, kidney, and heart and were placed directly into Trizol, homogenized and then snap frozen in liquid nitrogen. At least after 24 hours these tissue samples were then processed via standard protocols to isolate RNA. New York Genome Center RNA sequencing libraries were prepared using the KAPA Hyper Library Preparation Kit + RiboErase, HMR (Roche) in accordance with manufacturer’s recommendations. Briefly, 50-200ng of Total RNA was used for ribosomal depletion and fragmentation. Depleted RNA underwent first and second-strand cDNA synthesis followed by adenylation, and ligation of unique dual indexed adapters. Libraries were amplified using 12 cycles of PCR and cleaned-up by magnetic bead purification. Final libraries were quantified using fluorescent-based assays including PicoGreen (Life Technologies) or Qubit Fluorometer (Invitrogen) and Fragment Analyzer (Advanced Analytics) and sequenced on a NovaSeq 6000 sequencer (v1 chemistry) with 2×150bp targeting 60M reads per sample.

##### RNA-seq of COVID-19 hamster tissue samples

RNA from hamster tissues was extracted using TRIzol Reagent (ThermoFisher Scientific), followed by overnight precipitation at -20 °C, and quantified using a NanoDrop (ThermoFisher Scientific). 1 mg of total RNA was enriched for polyadenylated RNA species and prepared for short-read next-generation sequencing using the TruSeq Stranded mRNA Library Prep Kit (Illumina) according to the manufacturer’s instructions. Sequencing libraries were sequenced on an Illumina NextSeq 500 platform. Fastq files were generated using bcl2fastq (Illumina) and aligned to the Syrian golden hamster genome (MesAur 1.0, ensembl) using the RNA-seq Alignment application (Basespace, Illumina). Golden Hamster ensembl genes were matched to homologous external gene names, human homolog ensembl genes, and human associated homolog gene names using BioMart (*4, 5*). OrthoFinder was used to generate orthologous human ensembl gene ids and gene names (*6*). After further filtering and quality control, R package edgeR (*7*) was used to calculate RPKM and Log2counts per million (CPM) matrices as well as perform differential expression analysis.

##### RNA-seq of COVID-19 murine tissue samples

RNA from inferior mouse lung lobes was extracted using TRIzol Reagent (ThermoFisher Scientific), followed by overnight precipitation at -20 °C, and quantified using a NanoDrop (ThermoFisher Scientific). Ribosomal RNA from 1000 ng total extracted RNA was depleted using a NEBNext rRNA Depletion Kit (Human/Mouse/Rat) (New England Biolabs Inc.). The remaining RNA was used to produce the sequencing libraries using the NEBNext Ultra II Directional RNA Library Prep Kit for Illumina (New England Biolabs Inc.) with AMPure XP (Beckman Coulter Life Sciences) for all bead cleanup steps. The libraries were sequenced on a NovaSeq 6000 System, using a NovaSeq 6000 SP Reagent Kit v1.5 (Illumina).

#### QUANTIFICATION AND STATISTICAL ANALYSIS

##### Analysis of nasopharyngeal swab RNA-seq data

The nasopharyngeal swab samples were analyzed comparing COVID-19 viral infection to the negative patients as previously described in Butler et al. (*1*)and the DESeq2 (*8*)was utilized to generate the differential expression data. Using all clinical samples there were a total of N = 216 COVID+ samples and 519 COVID negative samples. There also included 17 positive (Vero E6 cells) and 33 negative (buffer) controls.

##### Analysis of autopsy RNA-seq data

The full methods for the analysis of autopsy patients can be found in the Park et al. (*2*). Briefly, RNA-seq data was processed through the nf-core/rnaseq pipeline (*9*). This workflow involved adapter trimming using Trim Galore! (https://github.com/FelixKrueger/TrimGalore), read alignment with STAR (*10*), gene quantification with Salmon (*11*), duplicate read marking with Picard MarkDuplicates (https://github.com/broadinstitute/picard), and transcript quantification with StringTie (*12*). Other quality control measures included RSeQC, Qualimap, and dupRadar. Alignment was performed using the GRCh38 build native to nf-core and annotation was performed using Gencode Human Release 33 (GRCH38.p13). FeatureCounts reads were normalized using variance-stabilizing transform (vst) in DESeq2 package in R for visualization purposes in log-scale (*8*). Differential expression of genes was calculated by DESeq2. Differential expression comparisons were performed as either COVID+ cases versus COVID- controls for each tissue specifically, correcting for sequencing batches with a covariate where applicable, or pairwise comparison of viral levels from the lung as determined by Counter data.

##### Metabolic modeling and downstream statistical analyses

We simulated the optimal fluxes of each metabolic reaction for each RNA-seq sample, using an adjusted version of a context specific constraint based metabolic modeling method (*13, 14*). We constructed a context specific genome scale metabolic model for each RNA-seq sample, by subsetting metabolic reactions in the Recon1 metabolic model (*15*) using CORDA (*16*) and cobrapy (*17*) based on the RNA-seq profiled expression levels of the genes associated with each metabolic reaction. While applying CORDA, we assigned high expression confidence score 3 to the top 35% highest expressed genes, medium confidence score 2 to the 35% to 85% highest expressed genes, and low confidence score 1 to 85%-100% highest expressed genes, in order to minimize the standard deviation of flux-levels across samples from each group for all groups. In addition, we manually activated the following essential pathways that are considered to be necessary for the model stability, since these essential pathways may not be properly activated based on RNA-seq data alone: ‘Oxidative Phosphorylation’, ‘Citric Acid Cycle’, ‘Glycolysis/Gluconeogenesis’, ‘CoA Biosynthesis’, ‘CoA Catabolism’, ‘NAD Metabolism’, ‘Fatty Acid Metabolism’, ‘Fatty acid activation’, ‘Fatty acid elongation’, ‘Fatty acid oxidation’, ‘ROS Detoxification’, and ‘Biomass and maintenance functions’. On each context specific genome scale metabolic model, we applied flux balance analysis (FBA) with each reaction as the objective to optimize the flux of the objective reaction. The LP optimization problems were solved by Gurobi solver (Gurobi Optimization, L. L. C., 2020) whose reference manual can be found in https://www.gurobi.com. All modeling procedures were implemented within a Jupyter notebook 6.1.5 using Python 3.7.7 (*18*). While gene expression levels were applied to the modeling procedure by each RNA-seq sample, all other parameters were maintained identically for all RNA-seq samples. The modeling procedure reported outcome as the FBA generated optimal flux levels of all available reactions of each context specific metabolic model constructed from each RNA-seq sample and the optimal flux levels were analyzed as variables for comparison between grouped cohorts as discussed in the text. Since it is impossible to presume variance or normality of flux distributions within and between cohort groups, a non-parametric Van der Waerden (VdW) test was applied to compare groupwise flux levels using the R matrixTests package (v. 0.1.9). Some pathways were chosen based on mitochondria-associated connection or Covid19-associated peculiarity and selected reactions with significant p values < 0.05.

##### Analysis of monocyte RNA-seq data

The monocyte COVID-19 RNA-seq data, published under the accession GSE159678 (*19*), was downloaded from SRA and gene expression was quantified using Salmon’s selective alignment approach (*20*). The RNA-seq processing pipeline was implemented using pyrpipe (*21*) (https://github.com/urmi-21/pyrpipe/tree/master/case_studies/Covid_RNA-Seq). Exploratory data analysis and differential expression analysis were performed using MetaOmGraph (*22*).

##### Analysis combining autopsy and nasopharyngeal swab RNA-seq data

To combine the results from the autopsy and nasopharyngeal swab RNA-seq data, we utilized the t-score values from the DESeq2 analysis. Heatmaps were displayed using pheatmap (*23*). Circular heatmaps were produced in R (version 4.1.0), using the Complex Heatmap (*24*) (version 2.9.4) and circlize (*25*) (version 0.4.12) packages.

##### Analysis of COVID-19 murine tissue RNA-seq data

From the RNA-seq data the reads were aligned to the *Mus musculus* BALB/c or C57B/6 genome (v1.100) and the SARS-CoV-2 genome MA10 (MT952602) using the CLC Genomics Workbench v20.0 (https://digitalinsights.qiagen.com/) with the RNA-seq and Small RNA Analysis pipeline and the RNA-Seq Analysis module with all standard settings, the read counts then calculated. From the pipeline the fold-change and p-adjusted values were calculated and utilized for all downstream analysis similar to the autopsy data. The murine RNA-seq data has been deposited in the NCBI BioProject database (https://www.ncbi.nlm.nih.gov/bioproject/) under the BioProject accession number PRJNA803057.

##### Gene Set Enrichment Analysis (GSEA)

For pathway analysis, we utilized fast Gene Set Enrichment Analysis (fGSEA) (*26*). Pathway analysis was done utilizing custom-made Gene Set files from either MitoPathway or the genes we curated available in the **Tables S2&S3**). Using fGSEA, all samples were compared to controls and the ranked list of genes was defined by the t-score statistics. The statistical significance was determined by 1000 permutations of the genesets (*27*).

##### Proteomic and transcriptomic analysis in the blood tissue and nasopharyngeal samples and CaCo-2 cells

The nasopharyngeal relative mRNA level change between severe and uninfected subjects. Changes in host protein levels are from published 24 hours post-infection CaCo-2 cells mass spectrometry-based proteomic data (*28*) (https://www.nature.com/articles/s41586-020-2332-7). 2D annotation enrichment for RNA-seq and proteomics data was performed using Perseus 1.6.14.0 for proteomic and transcriptomic analysis of the mitochondrial genes (*29*) (https://www.nature.com/articles/nmeth.3901 and (*30*) https://bmcbioinformatics.biomedcentral.com/articles/10.1186/1471-2105-13-S16-S12).

We downloaded whole blood transcriptome data and plasma proteome data from The COVIDome Explorer Researcher Portal (*31*). We used the following filters for transcriptome data: Category “Effect of COVID-19 status”, Platform “Blood”, Statistical test “Student’s t-test”, Adjustment method “none”, Sex “male” and “female”, Age Group “All”. We used similar filters for proteome data: Category “Effect of COVID-19 status”, Platform “SOMAscan”, Statistical test “Student’s t-test”, Adjustment method “none”, Sex “male” and “female”, Age Group “All”. We analyzed mitochondrial genes at both transcriptome and proteome levels and visualized the data using RStudio Desktop 1.3.1093 (*32*), ggplot2 version 3.3.2 and ggrepel version 0.8.2 ggrepel (*33*).

---

**Fig. S1.**
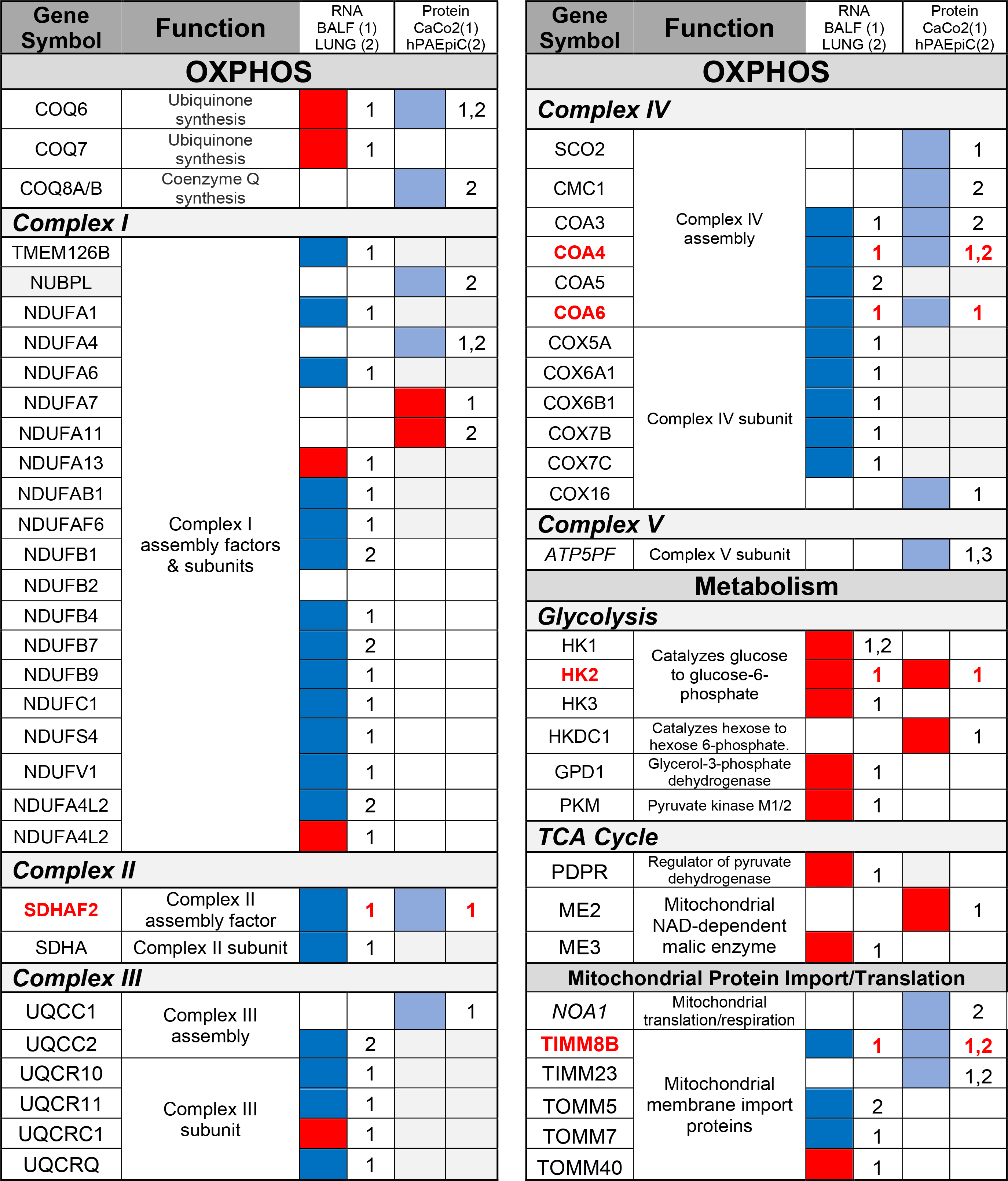
Conserved protein and transcript level changes in SARS-CoV-2 infection cells and tissues. We compared changes in host protein levels from infected CaCo-2 cells extracted from Bojkova *et al.* 2020 (*28*) (Protein 1) and human pulmonary alveolar epithelial cells (HPAEpiC) extracted from Wang *et al.* 2020 (*34*)(Protein 2) with changes in transcript levels from SARS-CoV-2 infected BALF (RNA 1) and lung autopsy (RNA 2) samples extracted from Miller *et al.* 2021(*35*). Blue squares represent decreased protein/RNA levels. Red squares represent increased protein/RNA levels. Bold red font represents conserved changes in both protein and transcript levels.

**Fig. S2.**
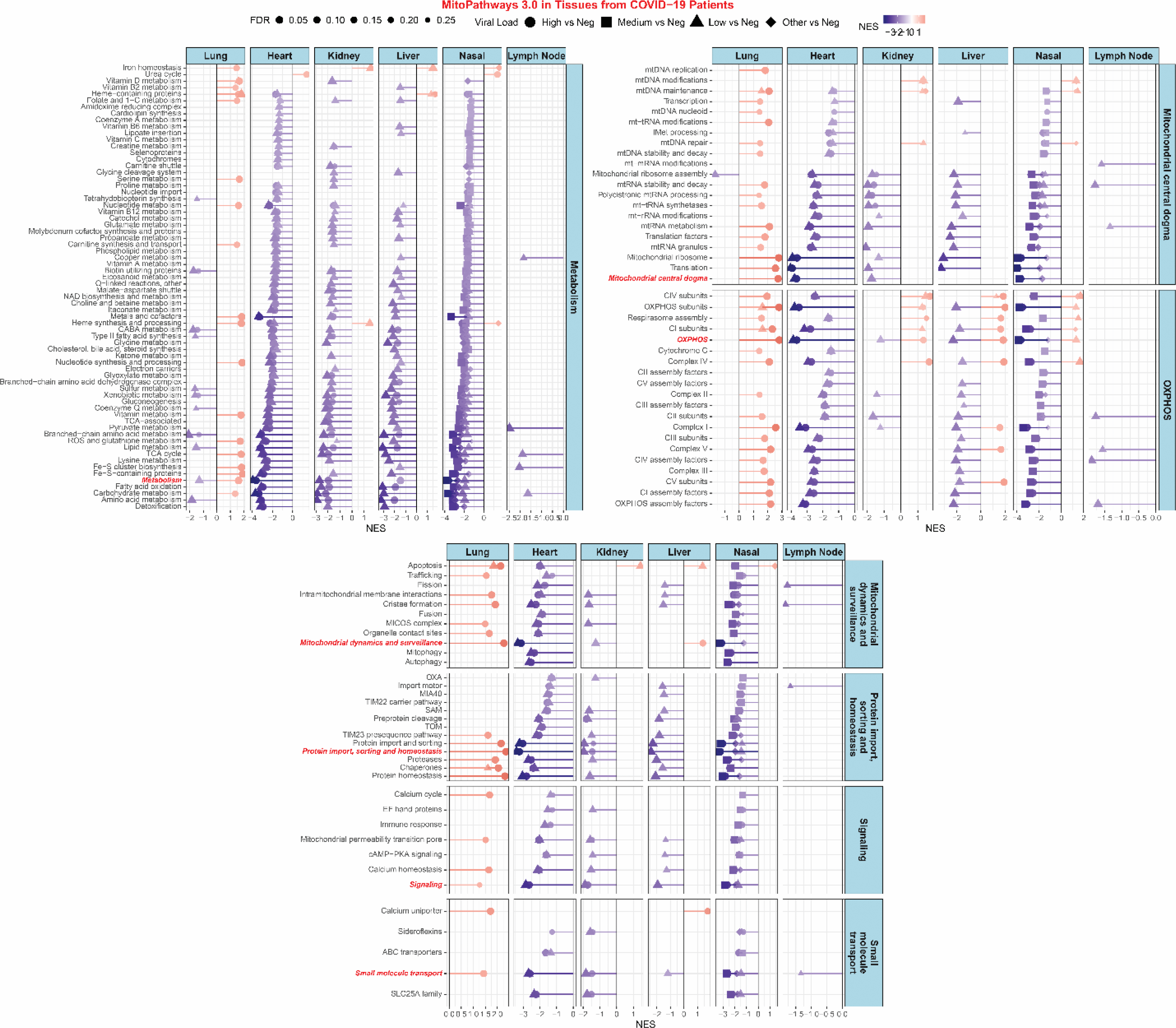
Lollipop plots for statistically significant MitoPathway gene sets determined by fGSEA. Only pathways with a FDR ≤ 0.25 are displayed. NES = nominal enrichment score.

**Fig. S3.**
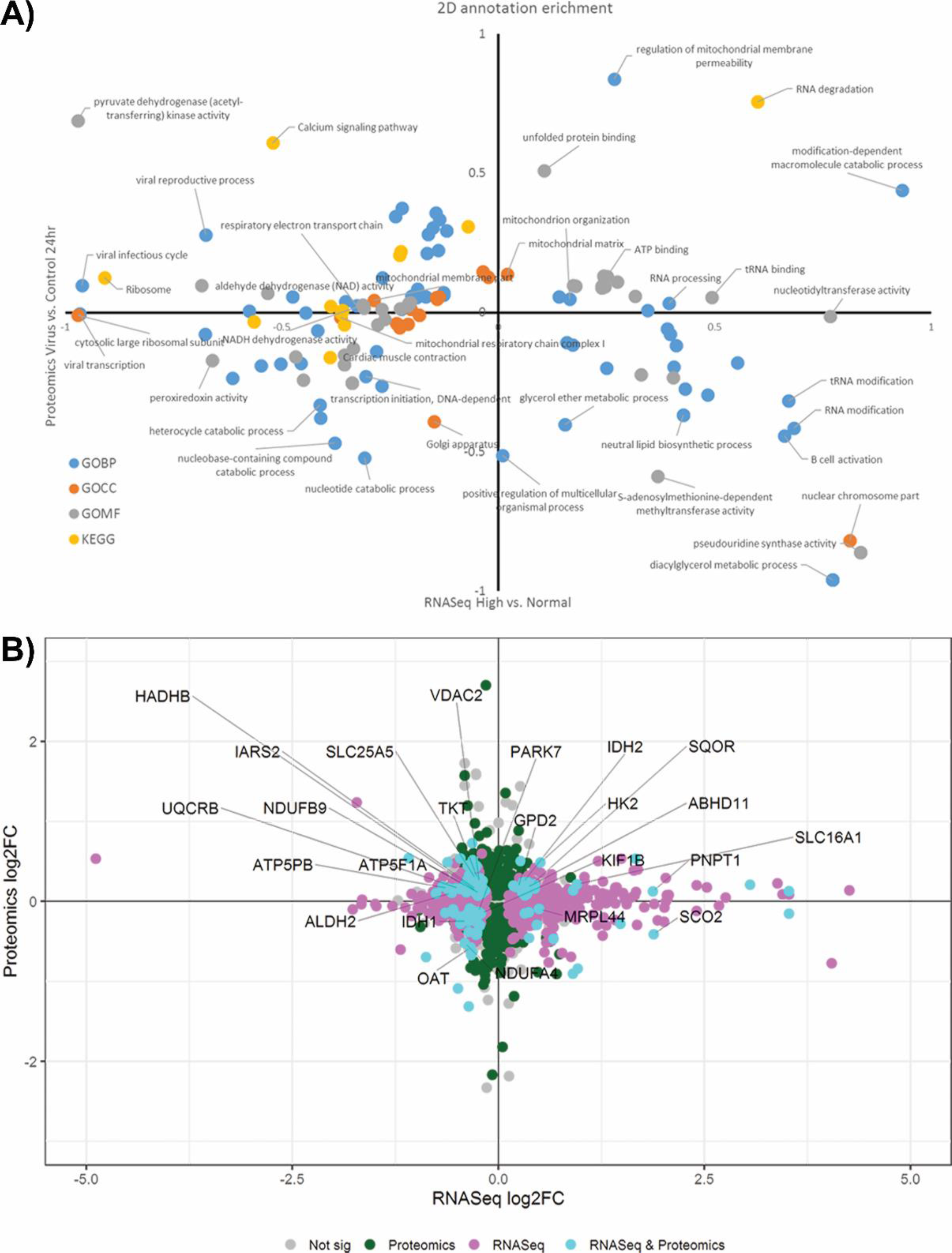
Correlation between mitochondrial protein and genes determined from MitoCarta. **A)** Gene Ontology (GO) analysis determined from nasopharyngeal RNA-seq patient data from high vs negative viral load compared to proteomic data from infected CaCo-2 cells (*28*). **B)** Mitochondrial genes overlapping between nasopharyngeal RNA-seq patient data from high vs negative viral load compared to proteomic data from infected CaCo-2 cells (*28*).

**Fig. S4.**
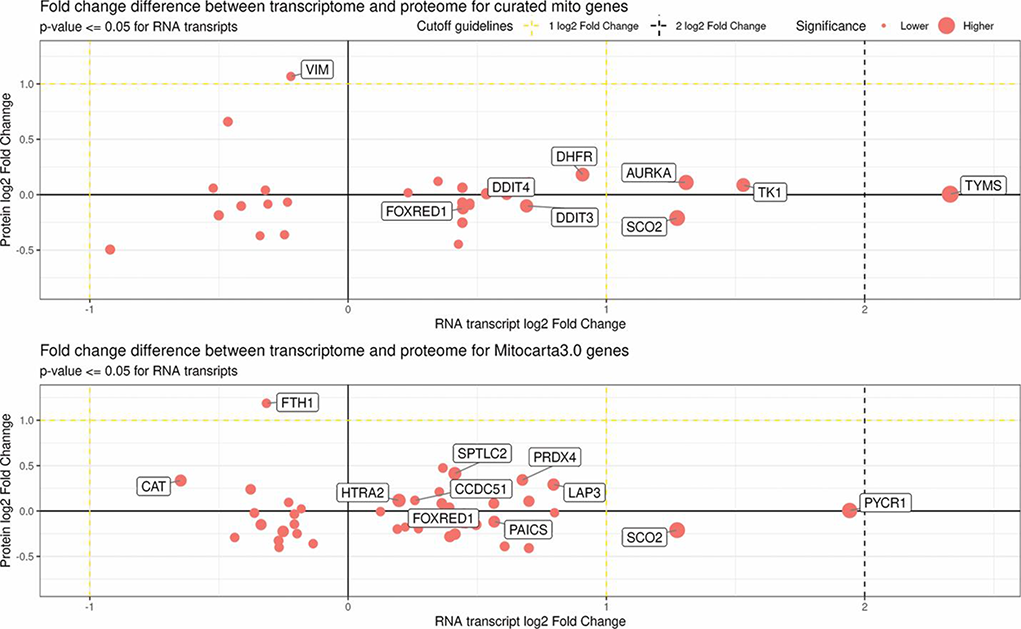
Correlation between mitochondrial protein and genes determined from MitoCarta in blood from COVID patients. Mitochondrial specific genes determined from our curated mitochondrial genes (top graph) and MitoCarta genes (bottom graph) comparing protein and RNA transcript changes in the blood from COVID patients (determined from COVIDome)(*31, 36*).

**Fig. S5.**
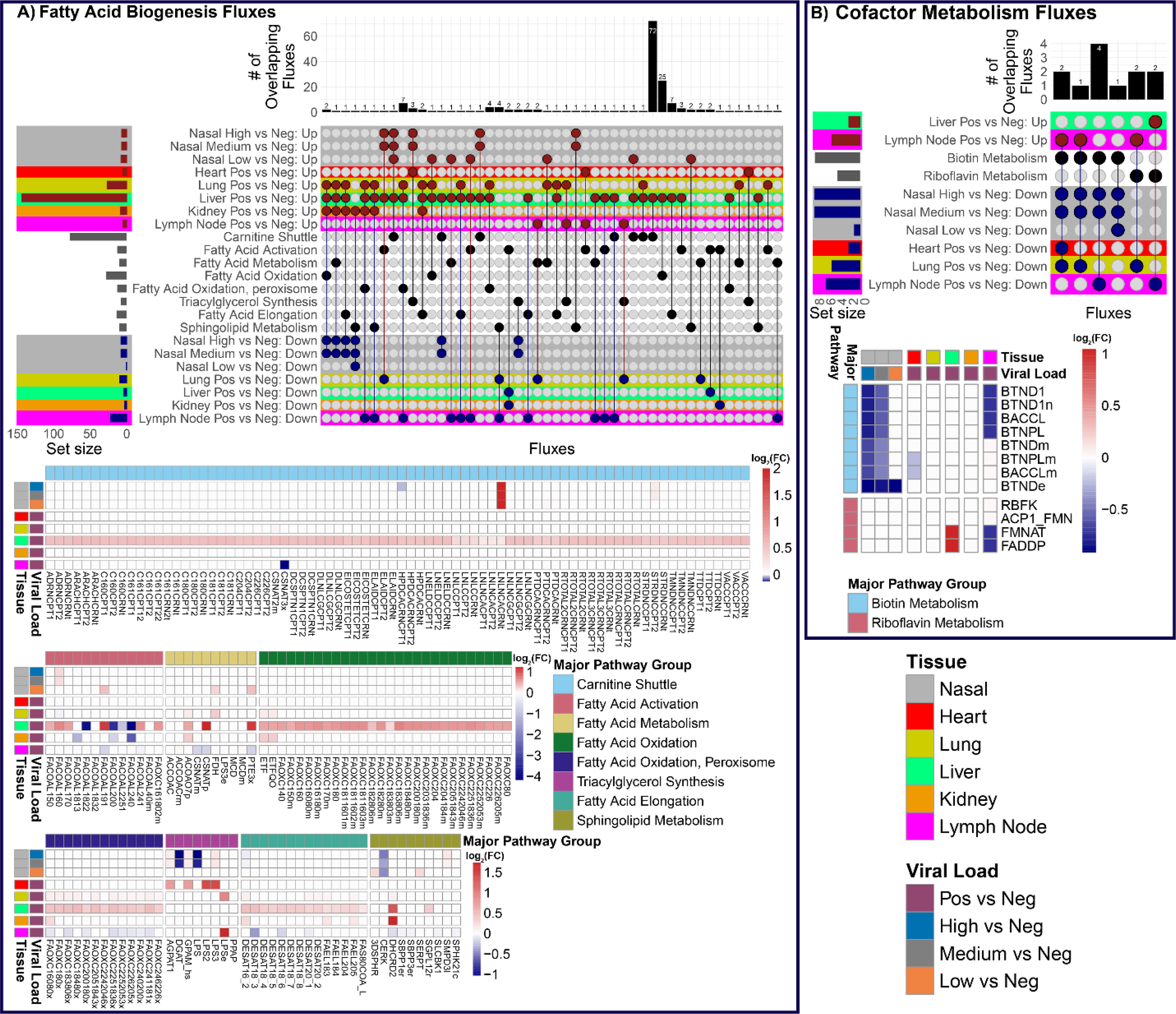
Metabolic flux analysis on RNA-seq data from nasopharyngeal and autopsy patient. A) Metabolic fluxes associated with fatty acid biogenesis. **B)** Metabolic fluxes associated with cofactor metabolism. For both **A) and B)** Upset plots are on the overlapping up- and down- regulation metabolic fluxes with the nasopharyngeal and autopsy RNA-seq patient samples. The red color dots represent the up regulated metabolic fluxes and the blue color dots represent the down regulated fluxes. The bar chart on top are the number of overlapping metabolic fluxes for each intersect. The set size bar plot represents the total number of metabolic fluxes contained in each row. Heatmaps of the of the log_2_ fold-change values for the significantly regulated fluxes in the major group flux pathways designated by the top or left color bar. The color bar represents the log_2_ fold-change values.

**Table S1.** Conserved Host-SARS-CoV-2 Mitochondrial Protein Interactions. We performed a comparative analysis of interactions between SARS-CoV-2 polypeptides and host mitochondrial proteins using data generated from three separate studies, Gordon et al. 2020 (37), Gordon et al. 2020 (38), and Stukalov et al. 2021 (39), via MitoCarta alignment. Jiang et al. 2020(40) and Wu et al. 2020 (41) are two additional studies that validated the Orf9b-TOMM70 interaction. Bold red font represents Host-SARS-CoV-2 interactions conserved between three or more studies.

| **Protein** | **Gene Symbol** | **Function** | **Citation** |
| --- | --- | --- | --- |
| ***Polypeptides of mitochondrial protein synthesis*** |  |  |  |
| M | TARS2 | Mitochondrial leucyl-tRNA synthetase | 37,38 |
|  | NARS2 | Mitochondrial asparaginyl-tRNA synthase | 39 |
| ***Polypeptides of OXPHOS*** |  |  |  |
| Nsp6 | ATP5MG | "g" subunit of the mitochondrial ATP synthase | 37,38 |
|  | ATP5F1E, ATP5F1B, ATP5PB, ATP5PD, ATP5F1D, MT-ATP6 | Subunits of the mitochondrial ATP synthase | 39 |
| M | **COQ8B** | **Coenzyme Q synthesis protein** | **37-39** |
| ***Polypeptides of the mitochondrial cytosolic protein import apparatus*** |  |  |  |
| M | PMPCA | Mitochondrial transit protease | 37,38 |
|  | **PMPCB** | **Mitochondrial transit protease** | **37-39** |
|  | **PITRM1** | **Mitochondrial pre-sequence protease** | **37-39** |
| Orf9b | **TOMM70** | **Outer mitochondrial membrane import protein** | **37-41** |
| Orf10 | TIMM8B | Inner mitochondrial membrane import protein | 37,38 |

**Table S2.**
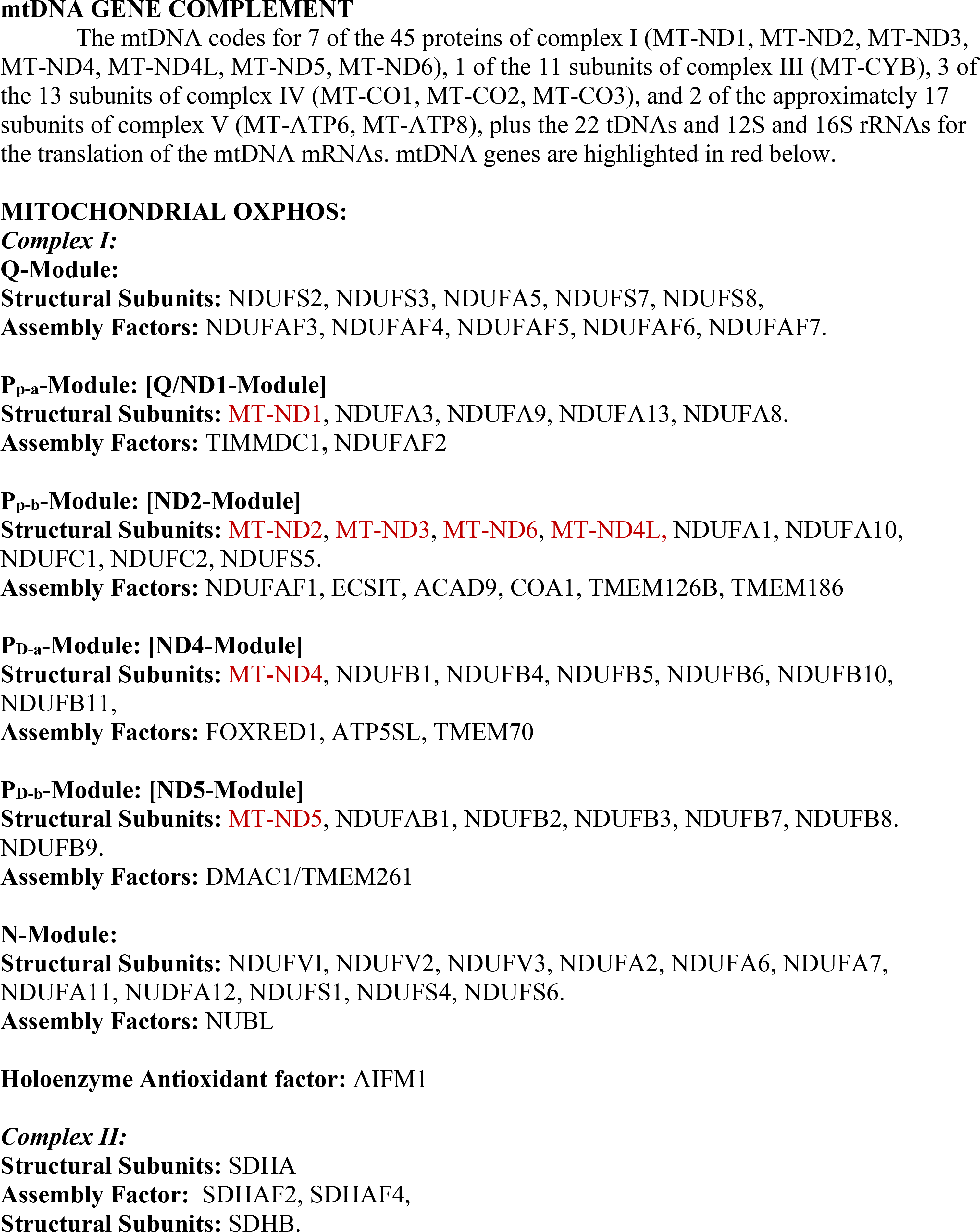
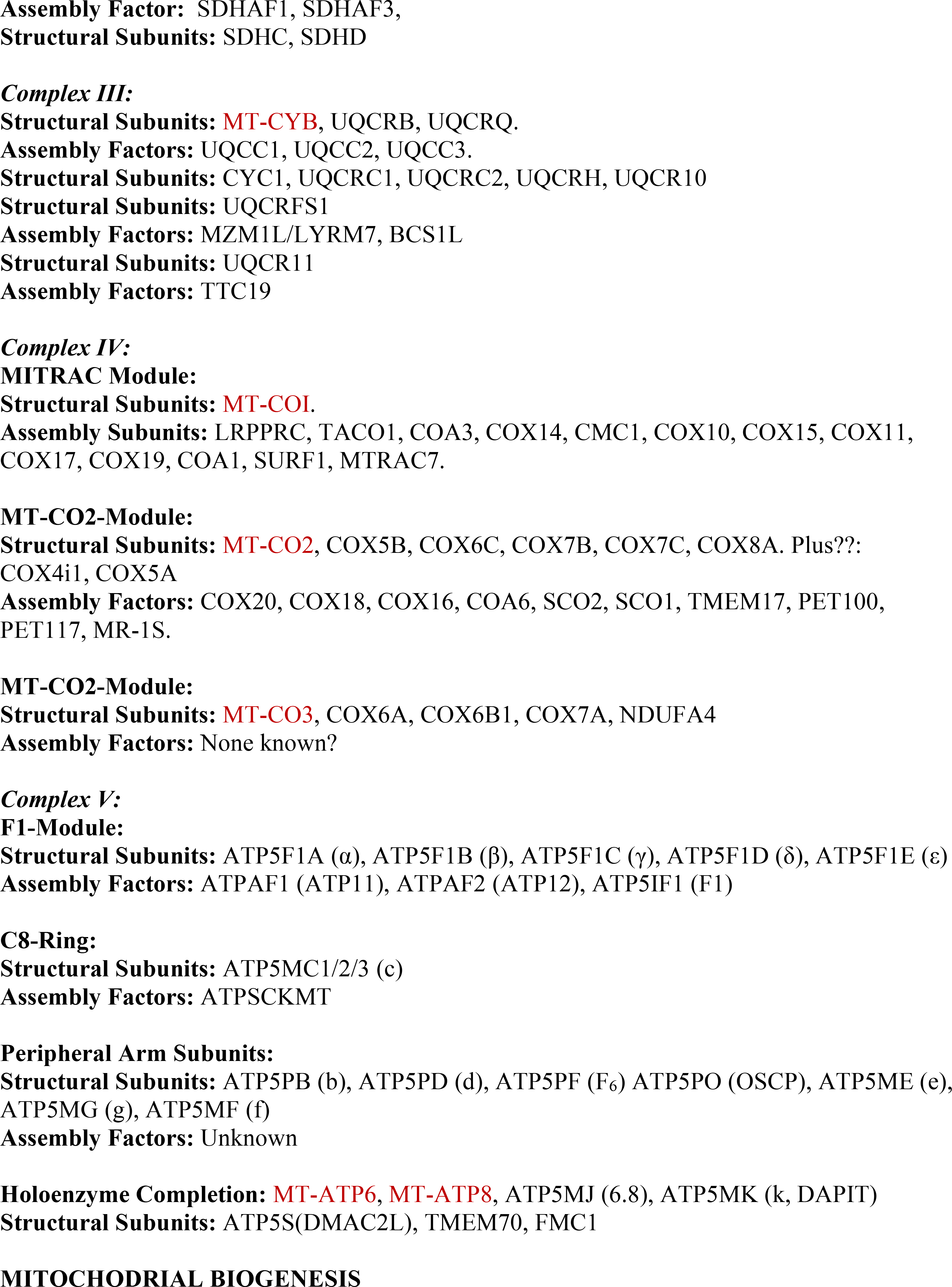
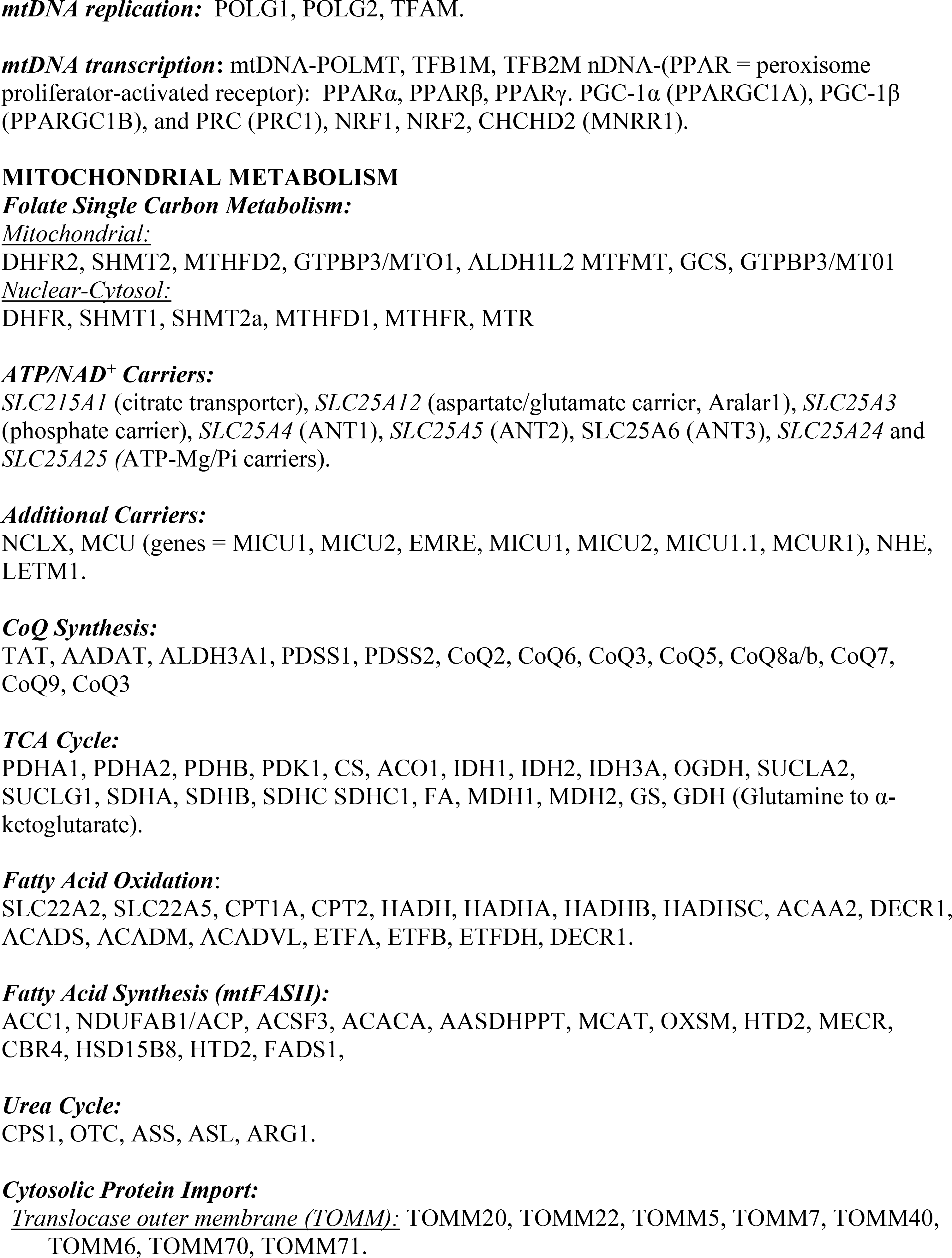
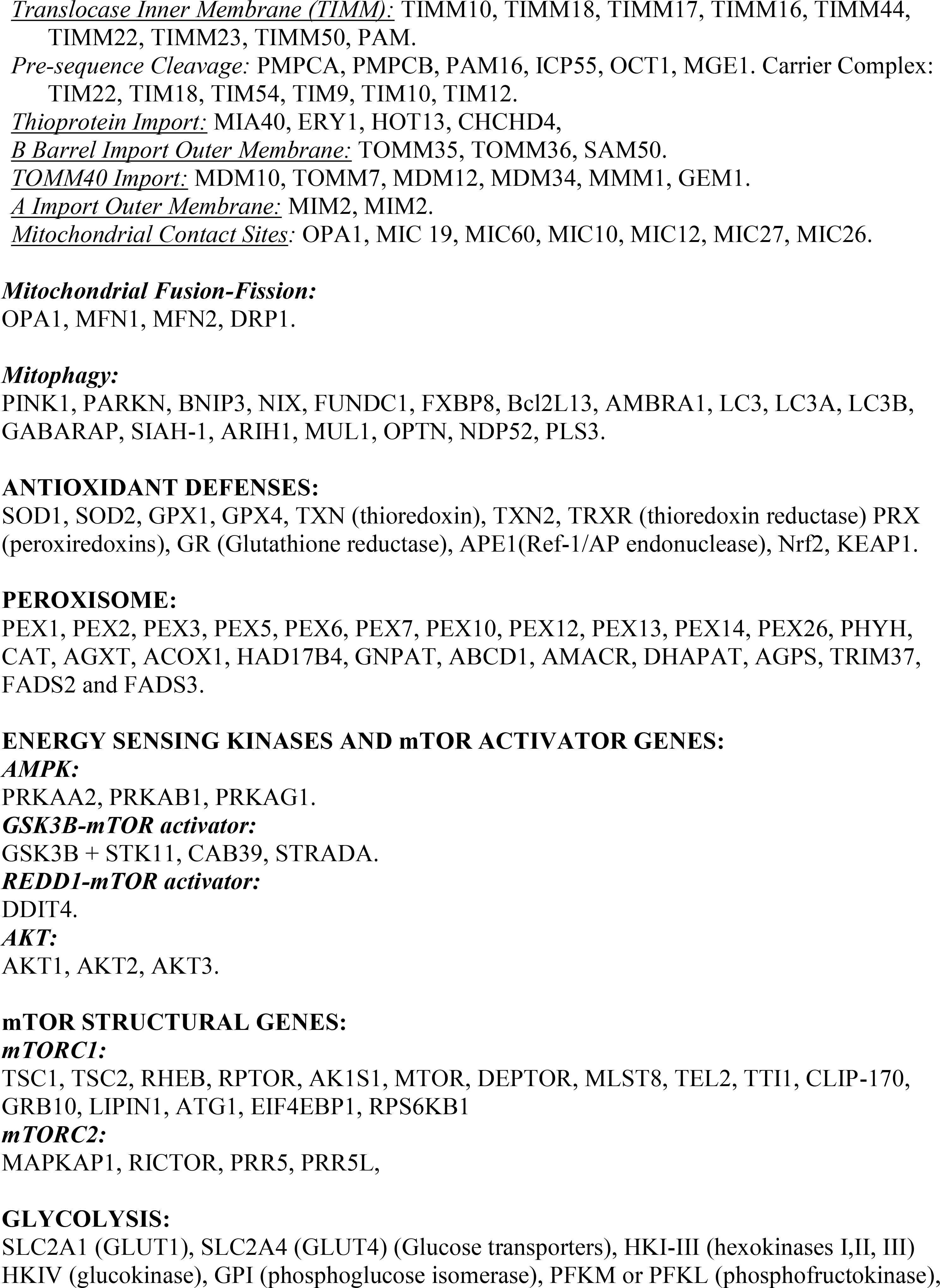
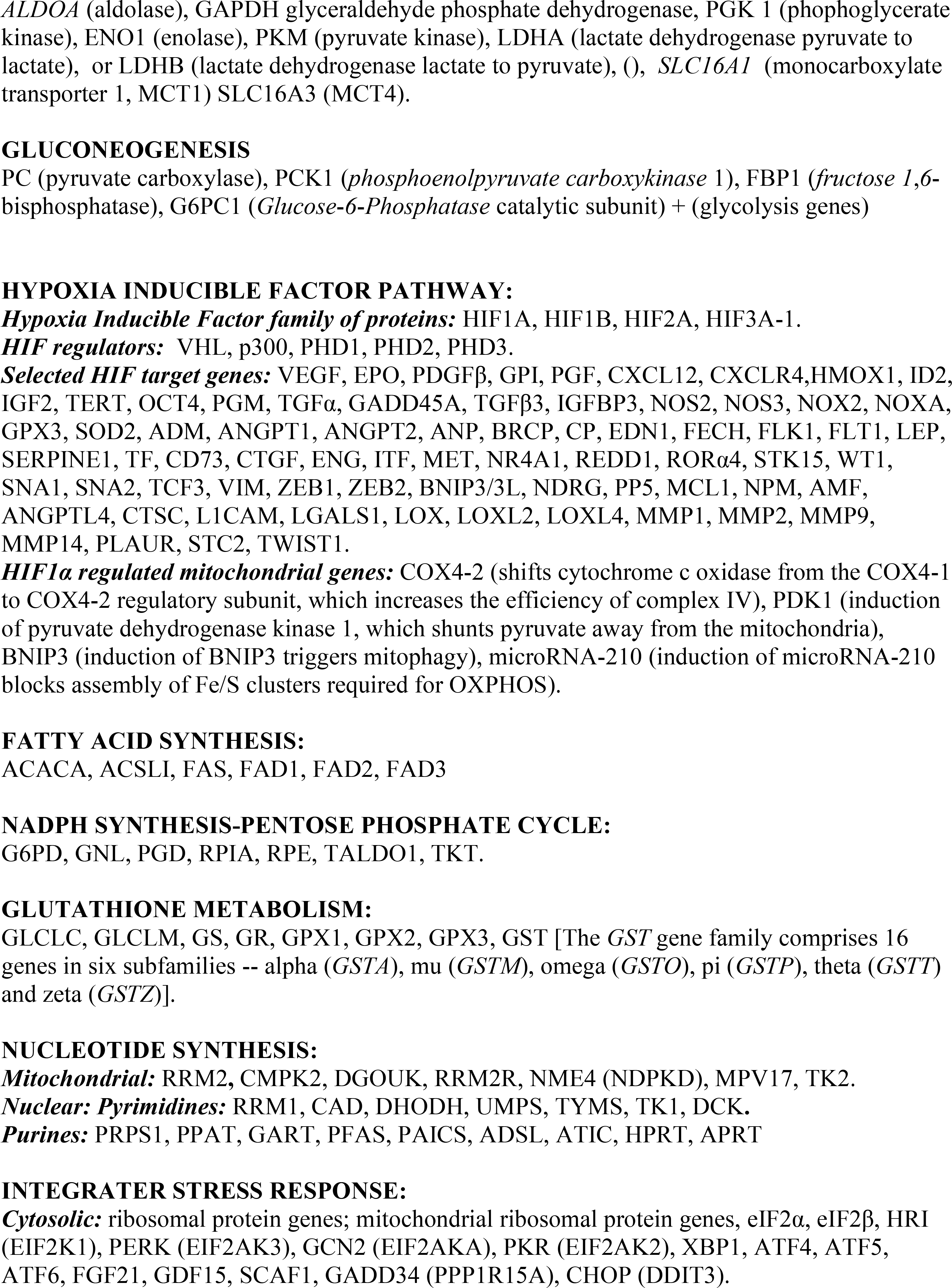
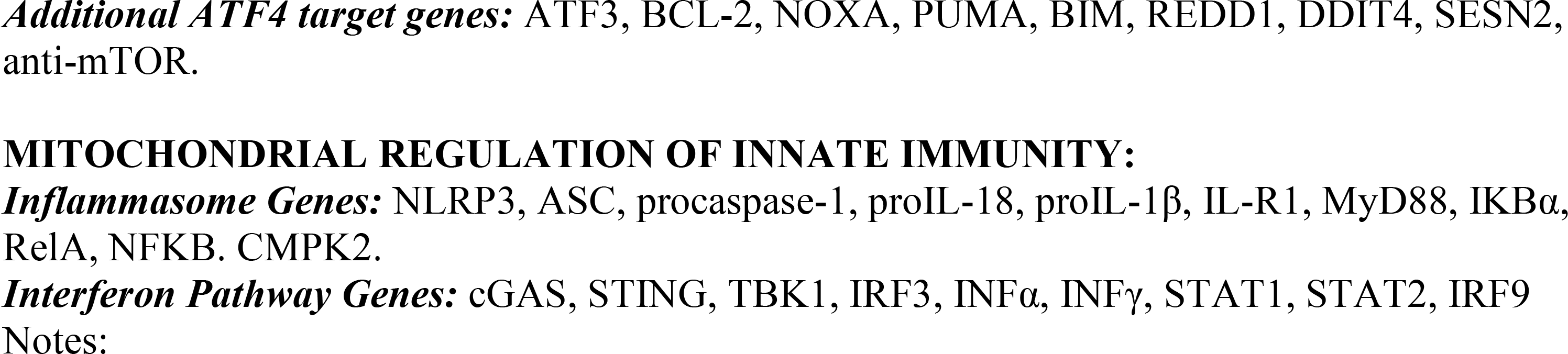
Curated mitochondrial genes utilized for Figs. 1 - 4.

**Table S3.**
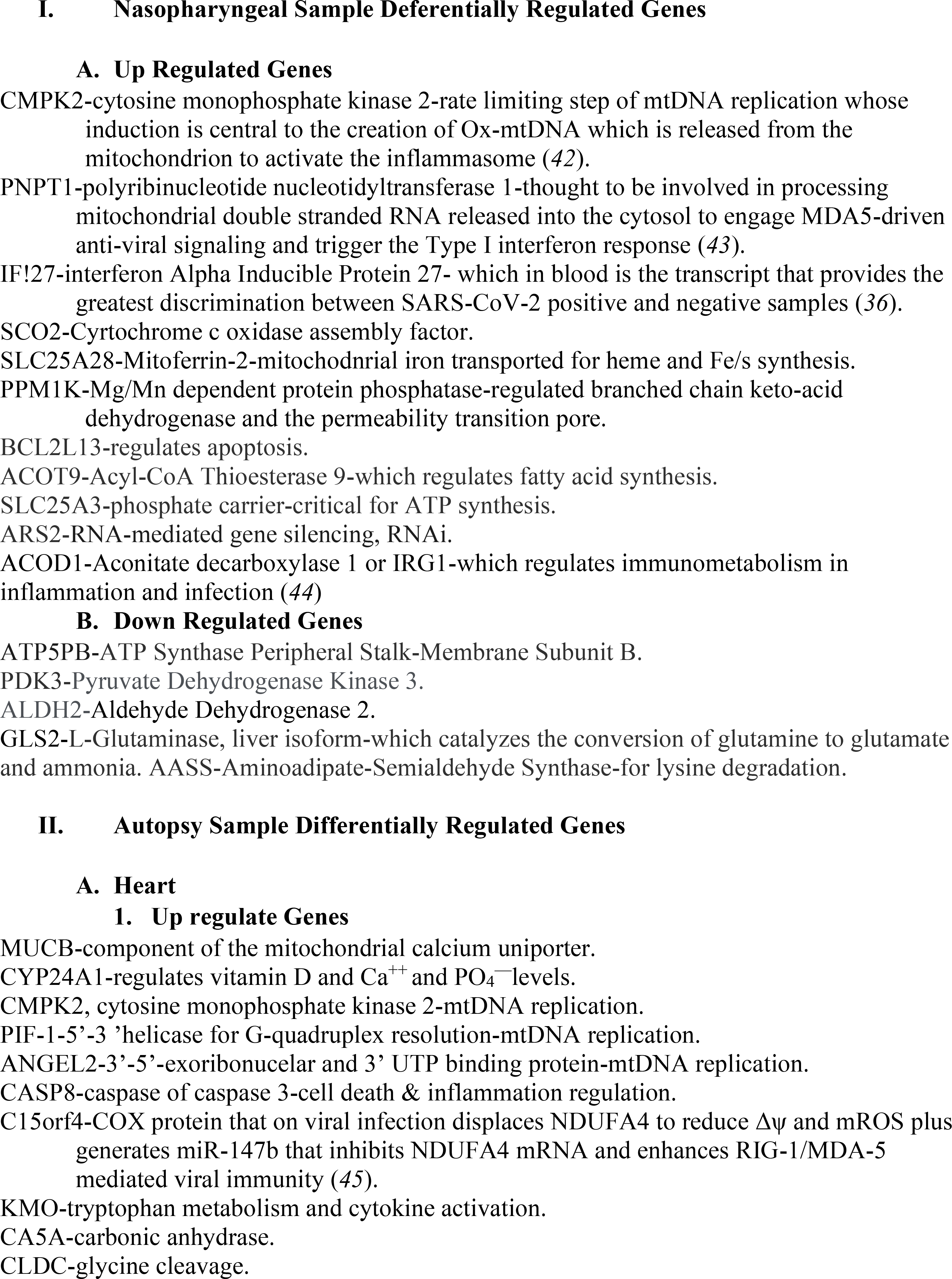
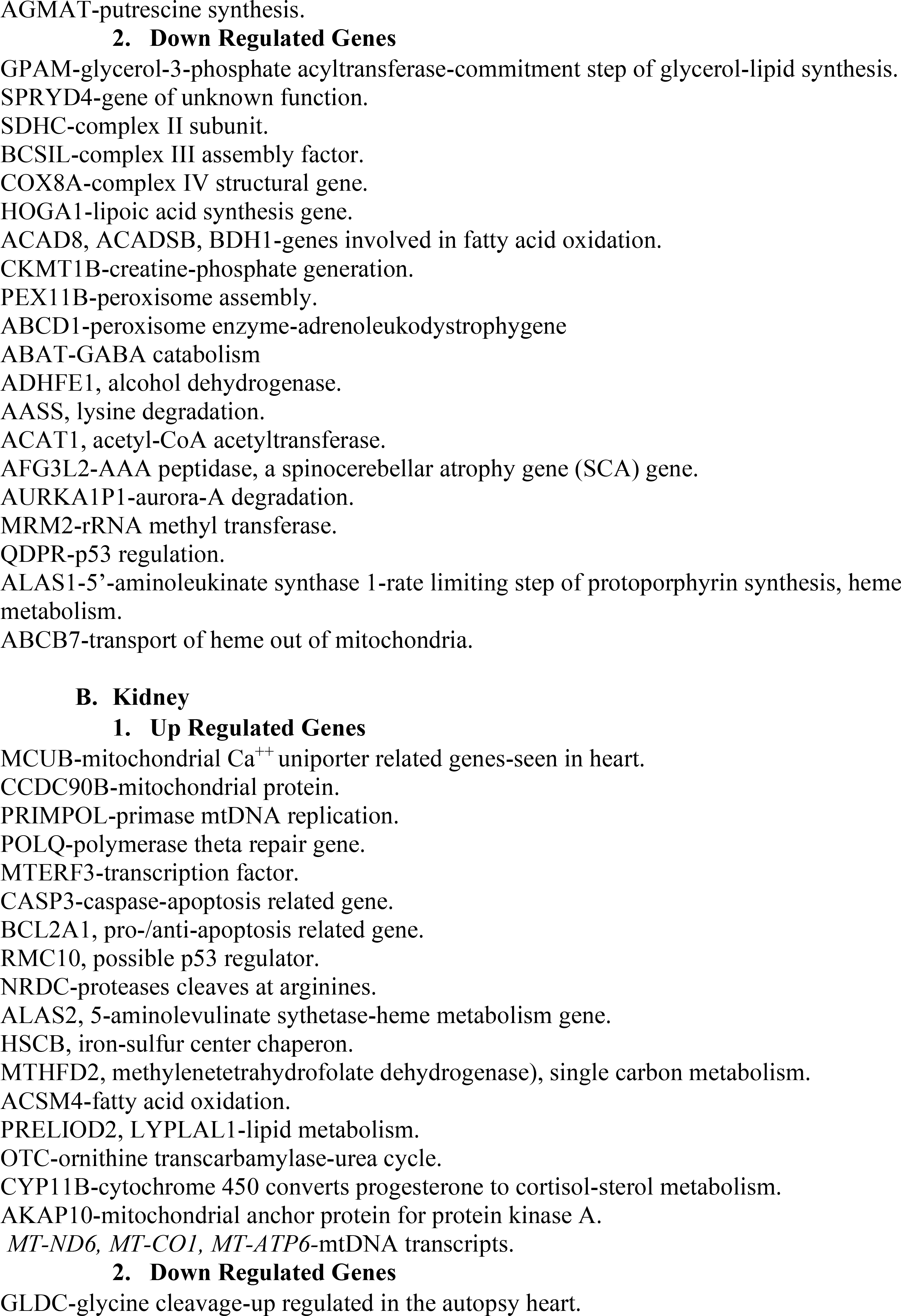
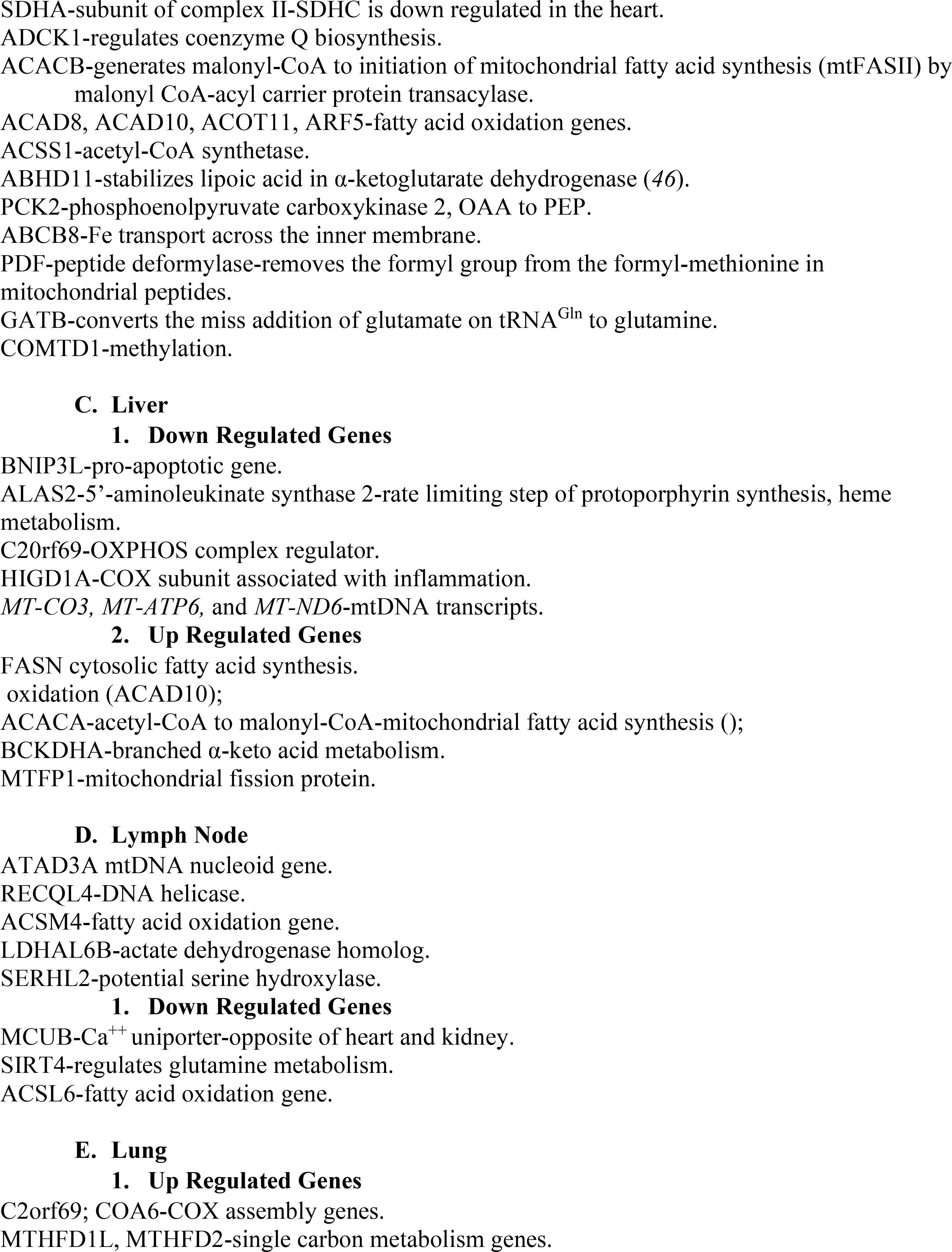
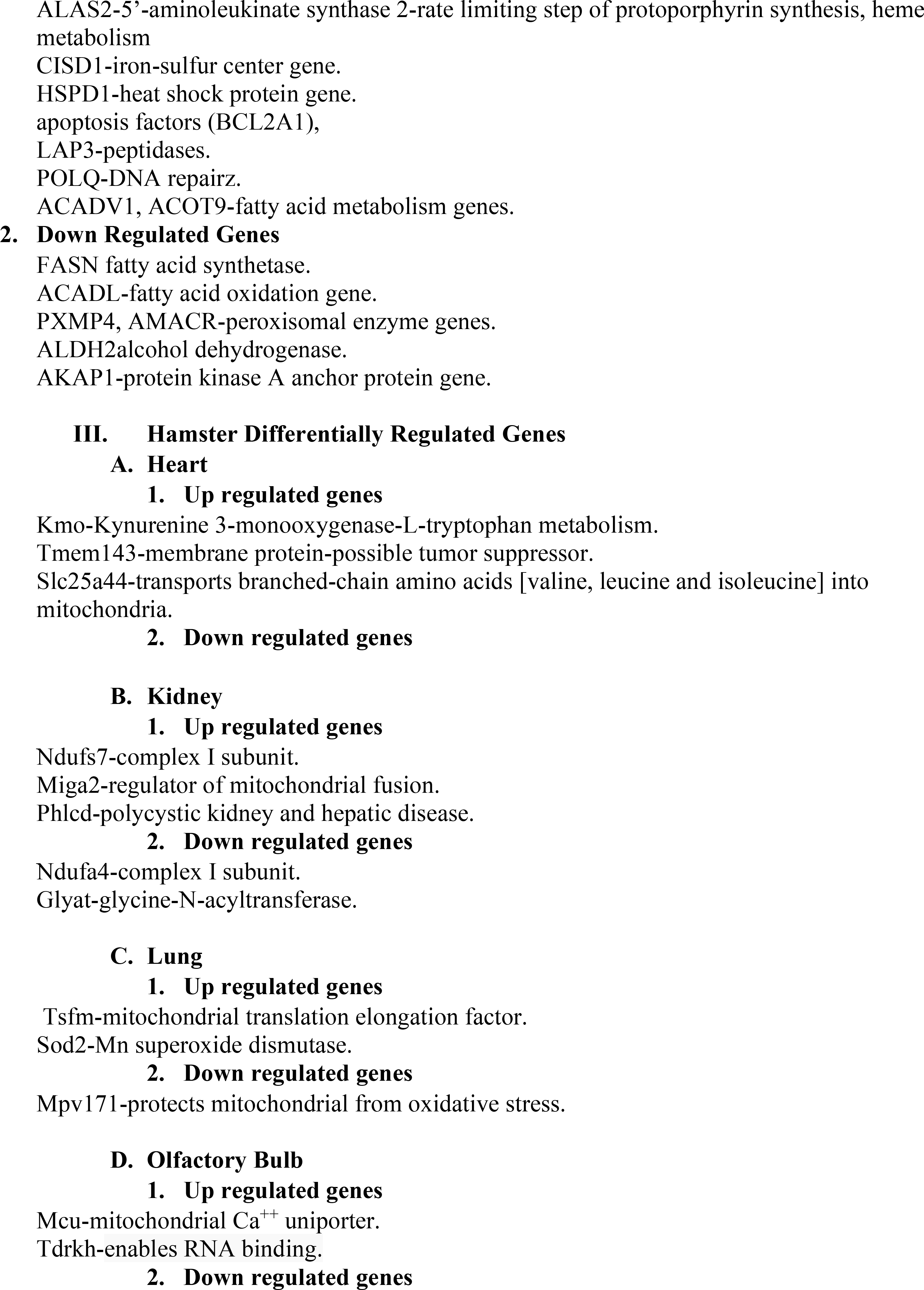
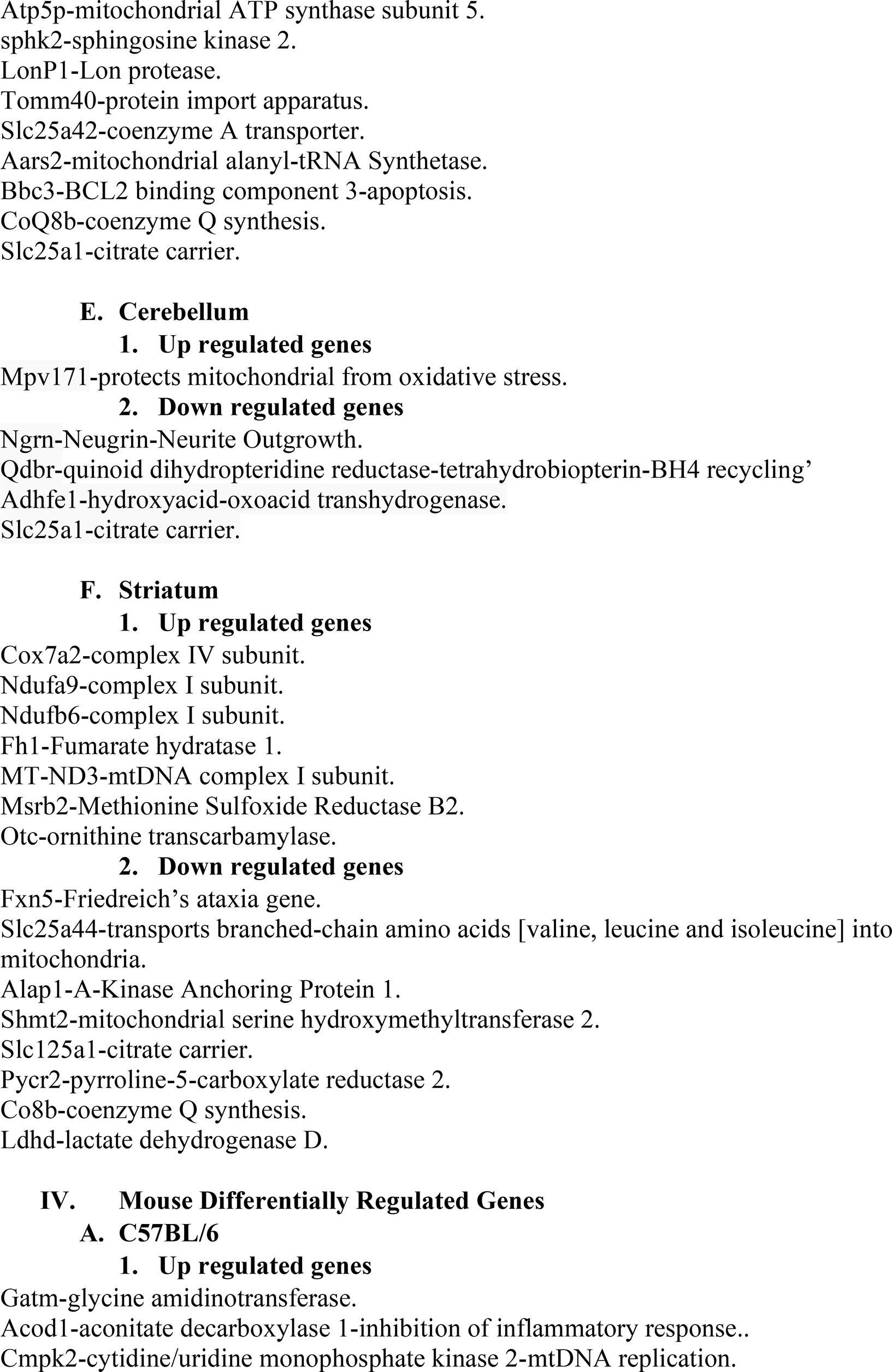
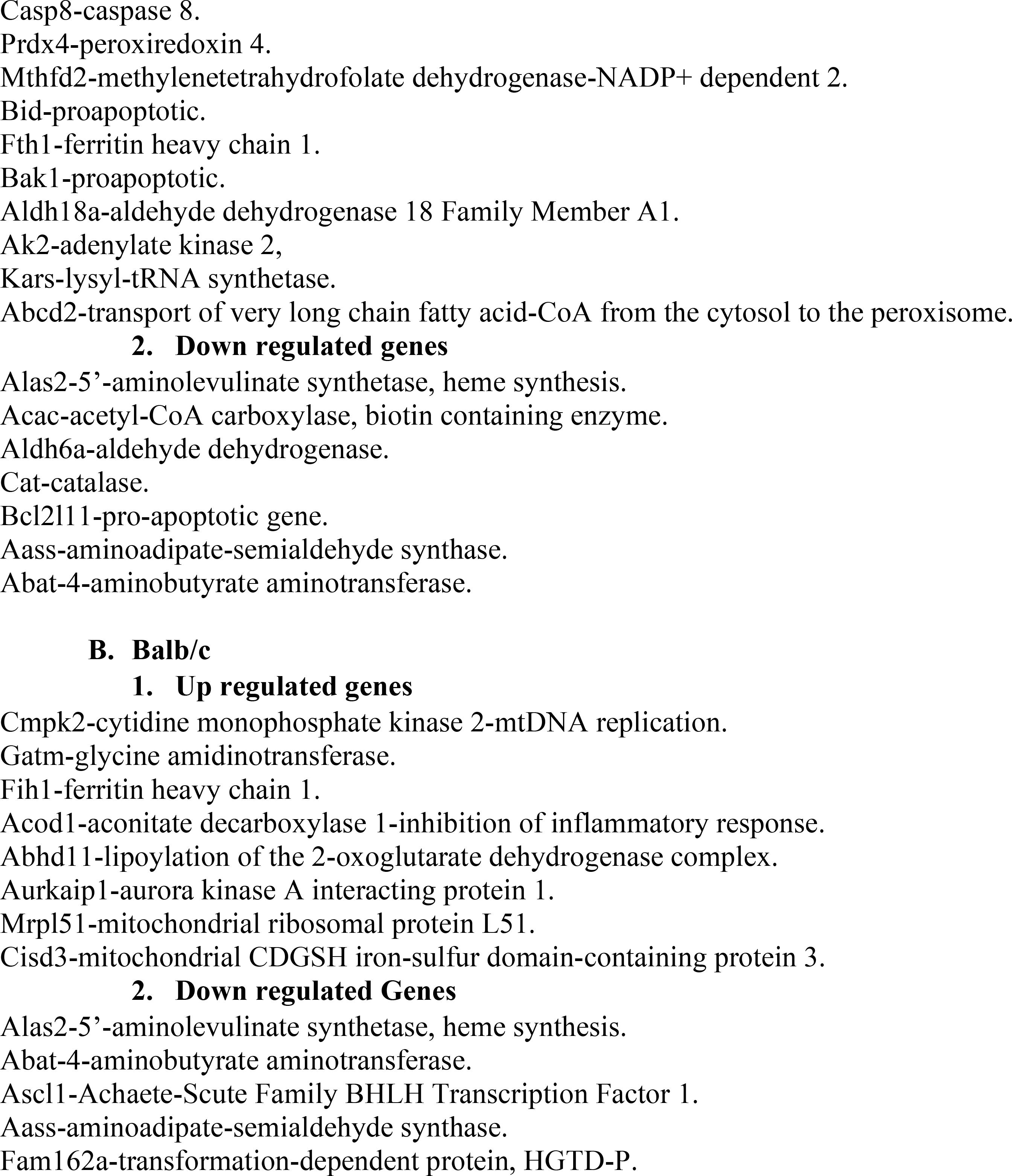
Representative Volcano Plot Genes that are Differentially Regulated.

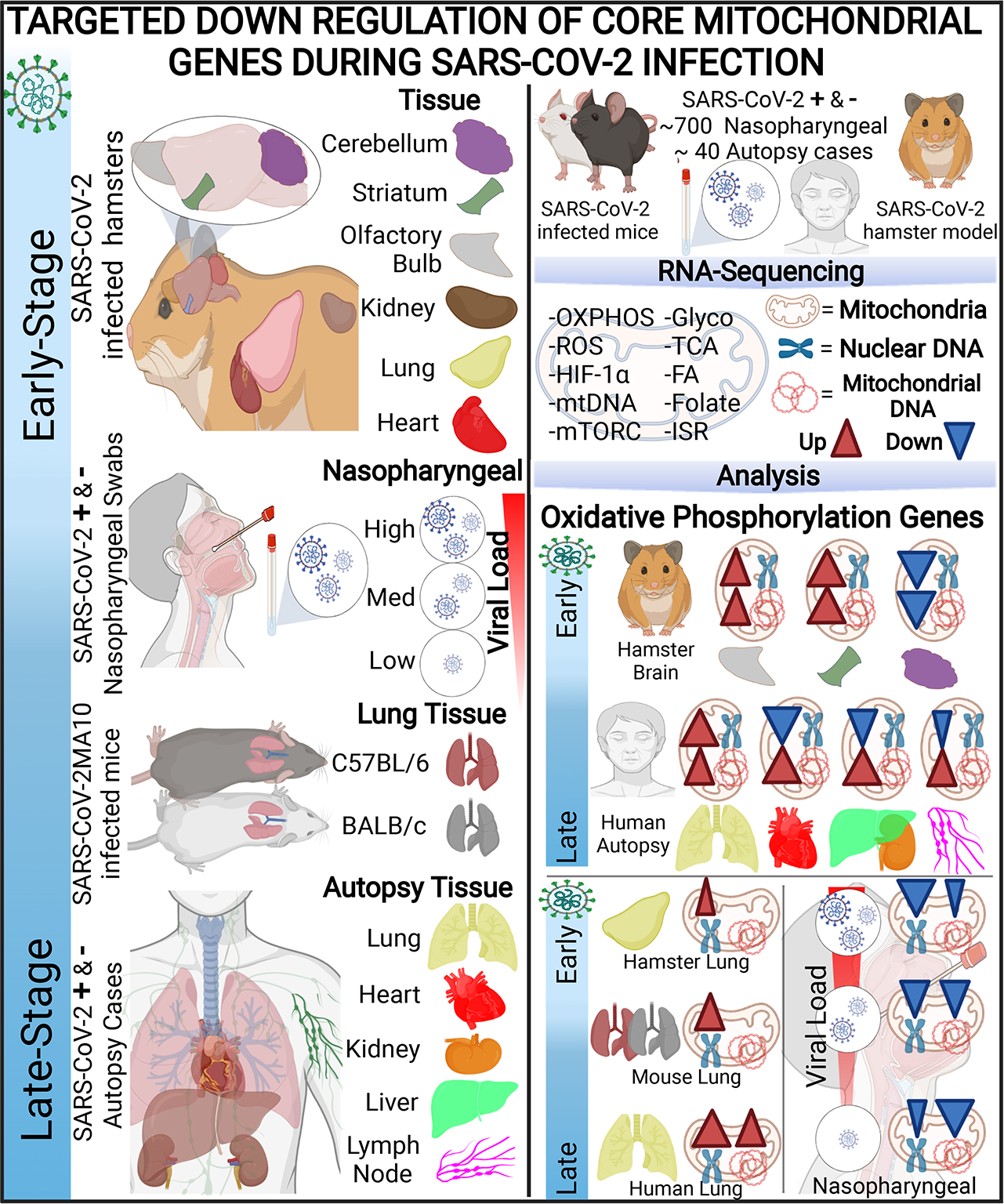

